# Mechanisms of GPCR hijacking by *Staphylococcus aureus*

**DOI:** 10.1101/2021.02.08.430206

**Authors:** Claire M. Grison, Cédric Leyrat, Paul Lambey, Sylvain Jeannot, Elise Del Nero, Simon Fontanel, Fanny Peysson, Joyce Heuninck, Rémy Sounier, Thierry Durroux, Sébastien Granier, Cherine Bechara

## Abstract

Atypical chemokine receptor 1 (ACKR1) is a G protein-coupled receptor (GPCR) targeted by *Staphylococcus aureus* (SA) bi-component pore-forming leukotoxins to promote bacterial growth and immune evasion. Here we have developed an integrative molecular pharmacology and structural biology approach in order to characterize the effect of leukotoxins HlgA and HlgB on ACKR1 structure and function. Interestingly, we found that both components HlgA and HlgB compete with endogenous chemokines through a direct binding to ACKR1 captured by native mass spectrometry (MS). Unexpectedly, HDX-MS analysis revealed that toxin binding allosterically modulates the intracellular G protein-binding domain of the receptor, resulting in dissociation of ACKR1–G protein complexes in living cells. Altogether, our study brings important molecular insights into the initial steps of leukotoxins targeting a host GPCR. Our findings may open the way to develop antibiotics inhibiting host receptors binding, a mechanism of action less prone to resistance.

## INTRODUCTION

*Staphylococcus aureus* (SA) is a major concern for public health mainly due to the emergence of various multidrug-resistant strains of this pathogen (Foster, 2004; Turner et al., 2019). SA produces a large arsenal of virulence factors, among which bi-component leukocidins -also referred to as leukotoxins -stand out as interesting targets for developing novel antivirulence strategies (Assis et al., 2017; Kong et al., 2016). Leukotoxins belong to the family of β-barrel pore-forming toxins that assemble into heteromeric pores to lyse specific cells (Alonzo and Torres, 2014; Los et al., 2013; Seilie and Bubeck Wardenburg, 2017; Vandenesch et al., 2012). Five different types of leukotoxins are expressed by SA infecting humans: PVL, LukED, HlgAB, HlgCB and LukAB. Each one is formed by two subunits, the host cell targeting S component (S for slow elution during biochemical purification) and the polymerization F (fast elution) component. In the current view, with the exception of LukAB, all the subunits are believed to be secreted as monomers that will heterodimerize upon specific interaction with host myeloid and erythroid cells. Dimerization will lead to a subsequent toxin oligomerization and pore-formation in cell membranes (Kaneko and Kamio, 2004; Yamashita et al., 2014, 2011).

Recent identification of leukotoxins receptors in targeted host cells increased our knowledge of the mechanism behind cellular specificity of these toxins and their role in pathogenesis (Alonzo III et al., 2012; Lubkin et al., 2019; Reyes-Robles et al., 2013; Spaan et al., 2014, 2015, 2017). Based on these findings, the predicted mechanism is that only the monomeric S component specifically interacts with various complement and chemokine receptors present on the surface of leukocytes, all related to the family of G protein-coupled receptors (GPCRs). The S-component later recruits the F-component to trigger oligomerization and pore formation. Though the F-components HlgB and LukD where shown to bind to the surface of erythrocytes independent from their S-component partners (Ozawa et al., 1995; Yoong and Torres, 2015), it was considered a receptor-independent binding. Recently, one of the F-components, LukF-PV, was shown to specifically require a receptor in order to recognize targeted cells (Tromp et al., 2018), challenging therefore the proposed initial steps of receptor recognition and pore formation (Tromp and van Strijp, 2020).

Out of all targeted receptors, the atypical chemokine receptor 1 (ACKR1, previously called DARC) (Horuk, 2015) recognised by both HlgA and LukE is a key player. Indeed, in addition to being expressed in myeloid cells, ACKR1 is expressed in erythrocytes and endothelial cells making it necessary for SA to escape our immune system, to grow and to cause cell death (Lubkin et al., 2019; Spaan et al., 2015). Unlike canonical chemokine receptors, ACKR1 lacks the conserved DRYLAIV motif and is thus structurally unable to activate G proteins upon chemokine engagement (Nibbs and Graham, 2013; Novitzky-Basso and Rot, 2012). Rather, it internalizes and transports chemokines to the degradative compartment, acting as a chemokine buffer by modulating chemokine concentration and bioavailability (Cancellieri et al., 2013; Hansell et al., 2011; Pruenster et al., 2009; Vacchini et al., 2016). The high-resolution structure of ACKR1 is still unknown, however its homologues of known structures share the highly conserved GPCR structure, consisting of a single polypeptide chain with three intracellular and extracellular loops, an external N-terminal region essential for the specificity of ligand binding, and an intracellular C-terminal region that is involved in receptor signalling. Although increasing amount of structural and molecular data of chemokine receptors are being discovered (Arimont et al., 2017; Kufareva et al., 2017), the structural immunology and pharmacology related to ACKR1 is still in its infancy.

Binding of leukotoxins to GPCRs is poorly understood at the molecular and structural level. Various residues in the loops of the rim domain of HlgA and LukE as well as a 4-residue region in the cap domain of HlgA were shown to be necessary for the haemolytic activity and/or binding to erythrocytes (Nariya and Kamio, 1997; Peng et al., 2018; Vasquez et al., 2020). From the receptor side, LukE and HlgA seem to target different regions of ACKR1 N-terminal part, a highly flexible region, whereas both require sulfation of tyrosine residues in this same part of the receptor (Spaan et al., 2015). In addition, little is known on the effects of leukotoxins on ACKR1 binding to its natural ligands and downstream molecular signalling. Although biochemical and cell biology work has been done since the discovery of receptors targeted by leukotoxins, direct evidence capturing purified leukotoxin–receptor complexes has only been provided for the LukE–CCR5 pair (Alonzo III et al., 2012).

In this study, we used an integrative molecular pharmacology and structural mass spectrometry (MS) approach in order to characterize the effect of HlgAB binding on ACKR1 structure and function. We demonstrate that both leukotoxins, HlgA and HlgB, form independent complexes with purified ACKR1 *in vitro* using native MS (nMS). In living cells, TR-FRET experiments revealed that both HlgA and HlgB binding to ACKR1 compete with CCL5, an endogenous ligand. We also monitored the effect of leukotoxin binding on ACKR1 conformation using a combination of Hydrogen/Deuterium exchange-MS (HDX-MS) and cell-based resonance energy transfer. Surprisingly, in addition to the expected accessibility changes in the extracellular domain of the receptor, binding of leukotoxins induced long-range allosteric conformational changes in the intracellular domain of ACKR1 that leads to the dissociation of preassembled ACKR1–G protein complexes. Altogether, our study brings novel insights into the initial steps of leukotoxins biology through GPCR, namely the toxins effect on GPCR structure and function.

## RESULTS AND DISCUSSION

### nMS reveals HlgB and HlgA homodimers with HlgB being more prone to dimerization

We first analysed purified recombinant HlgA and HlgB by nMS in order to verify their oligomeric state, in the absence and the presence of detergent micelles as a mimic of the amphiphilic membrane environment. nMS has been gaining ground and has become a key actor in studying membrane protein interactions and dynamics (Allison and Bechara, 2019; Bechara and Robinson, 2015; Calabrese and Radford, 2018; Engen et al., 2020; Keener et al., 2020; Martens and Politis, 2020). It preserves non-covalent interactions and gives information regarding the stoichiometry and binding partners of protein complexes, among which GPCRs (Yen et al., 2017, 2018).

Surprisingly, HlgA and HlgB are present in a monomeric and homodimeric form in aqueous buffer (Fig. S1A), even at the lowest concentration analysed (sub-5 μM). The presence of detergent in the buffer enhanced the dimerization of both leukotoxins, as we detected an increase in the dimer-to-monomer ratio in the presence of DDM (Fig. 1A and B). Indeed HlgA and HlgB were able to bind multiple DDM molecules in their monomeric and dimeric forms (Fig. S1B), and detergent molecules were easily dissociated upon higher activation in the gas phase. Finally, an equimolar mixture of HlgA and HlgB resulted in the formation of heterodimers *in vitro*, in the absence of any receptor (Fig. 1C). In all analysed conditions, HlgB was systematically more prone to homodimerization than HlgA, since the dimer-to-monomer ratios at a given concentration was always 2-fold times higher for HlgB compared to HlgA. Taken together, our results demonstrate the presence of toxin homo- and hetero-dimers in solution even in the absence of membranes and receptors, and that detergent micelles enhances leukotoxin dimerization.

**Figure 1.**
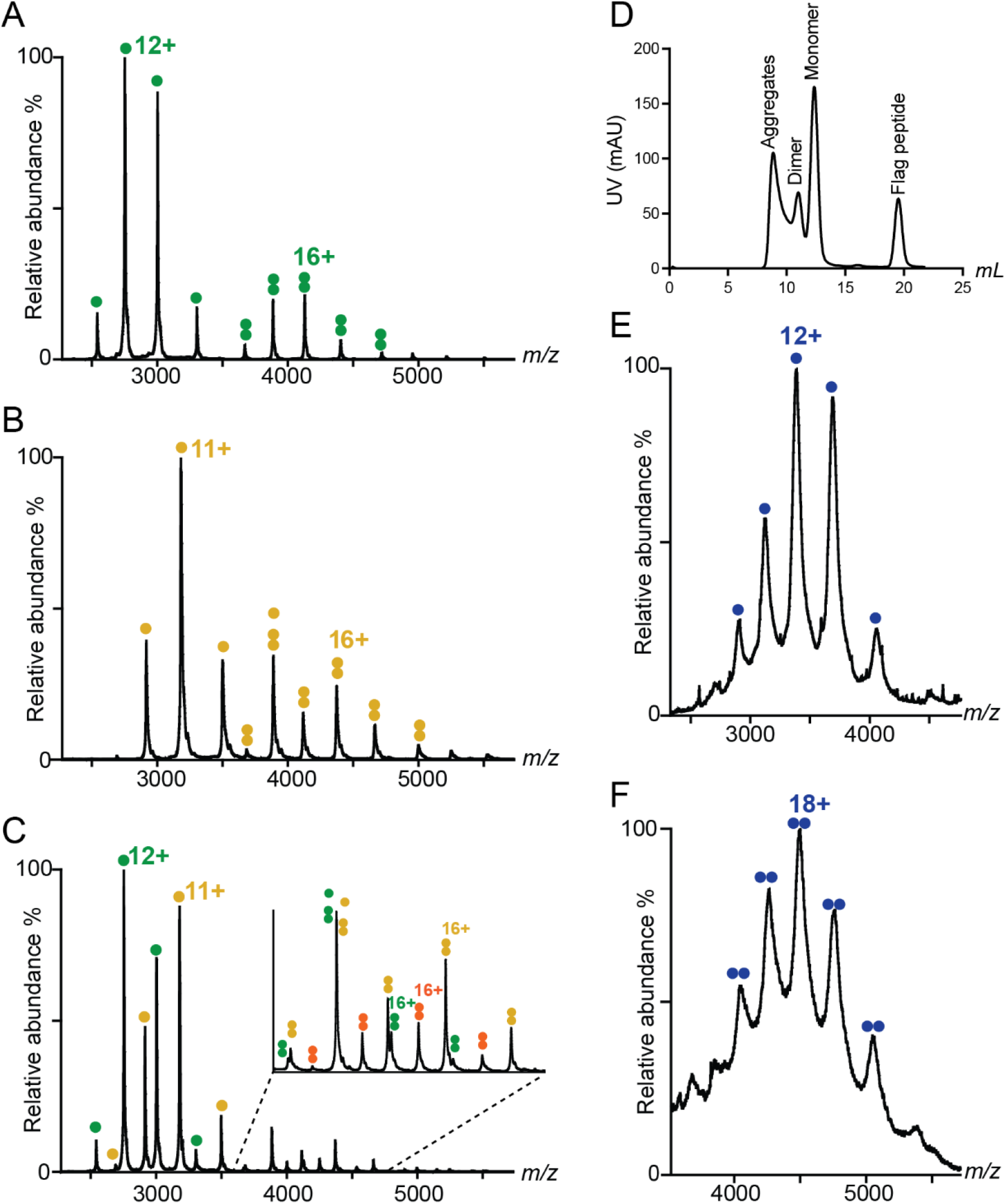
Native MS spectra of leukotoxins and ACKR1. A) nMS spectrum of 30 µM HlgA and B) 15 µM HlgB showing the presence of both monomeric (33 004 ± 1 Da and 34 943 ± 1 Da respectively) and dimeric (66 061 ± 14 Da and 69 972 ± 10 Da respectively) species. C) nMS spectrum of a mixture of 10 μM HlgA and 10 μM HlgB showing the additional presence of HlgAB heterodimers (69 925 ± 9 Da). Relative quantification in this equimolar mixture clearly shows more homodimeric HlgB compared to heterodimeric HlgAB and very small amount of homodimeric HlgA: 100% HlgA, 70.3% HlgB, 1.5% 2HlgA, 9.5% HlgAB and 27.7% 2HlgB. D) SEC profile of purified ACKR1. E) nMS spectrum of monomeric and F) dimeric ACKR1 produced in HEK GnTI^-^ cells (40 531 ± 32 Da and 81 032 ± 50 Da respectively) showing an additional ∼3.5 kDa glycosylations per monomer. All samples were buffer exchanged in 200 mM ammonium acetate pH 7.4 supplemented with 2CMC DDM prior to nMS analysis. Green circles HlgA, yellow circles HlgB, orange double circles HlgAB, blue circles ACKR1 WT.

### Purification of homogeneous human atypical chemokine receptor ACKR1

The GPCR ACKR1 was first produced in Sf9 cells and purified in DDM detergent, which gave a homogeneous monodisperse main peak by size-exclusion chromatography, as well as an additional smaller peak eluting at a higher apparent mass (Fig. 1D). nMS analysis of both peaks shows the presence of both monomeric and homodimeric ACKR1, however with additional heterogeneous masses around 3.5-5 kDa per monomer (Fig. S2A). In order to identify the nature of the observed additional mass, we produced ACKR1 in a cell line with restricted and homogeneous N-glycosylations (HEK GnTI^-^), which decreased the heterogeneity but did not lead to the complete removal of ACKR1 modifications (Fig. 1E, F and Fig. S2B). Finally, treatment of purified receptor with PNGaseF resulted in a complete removal of the modification, implicating that they indeed are N-glycosylations (Fig. S2C). We then verified all reported potential N-glycosylation sites by point mutations and found that all three N16, N27 and N33 are glycosylated. Mutating all glycosylation sites however resulted in receptor with a less stable Nter region, most probably due to the long and unstructured nature of the latter, since Nter hydrolysis products with purified N^16,27,33^Q-ACKR1 mutant were observed (Fig. S2D).

### HlgB and HlgA both bind separately to monomeric ACKR1 but not concomitantly

In order to provide a direct evidence of leukotoxin–ACKR1 binding and determine the stoichiometry of this complex, we analysed mixtures of purified recombinant HlgA, HlgB and monomeric ACKR1 using nMS. Incubating ACKR1: HlgA in a 2:1 ratio, ACKR1 being used at higher concentration to overcome lower ionization efficiency, for 30 min at 4°C revealed the presence of m/z species with a mass corresponding to proteins alone, as well as the presence of an ACKR1–HlgA complex with a 1 to 1 stoichiometry (Fig. 2A). Binding of HlgA gave similar results with WT and deglycosylated ACKR1, with peaks better defined in the mass spectra in the latter case. These results thus evidence the existence of leukotoxin–GPCR complex in detergent micelles and suggest that receptor glycosylation is not strictly necessary for ACKR1–HlgA interaction to occur. Surprisingly, we detected the formation of ACKR1– HlgB complexes when mixing the F-component HlgB with the receptor (Fig. S3), demonstrating a specific interaction between HlgB and ACKR1. Finally, when mixing the three components ACKR1: HlgA: HlgB in a 2: 1: 1 ratio, we detected separately formed ACKR1–HlgA and ACKR1–HlgB complexes, but no ternary ACKR1–HlgA–HlgB complexes were visible. Taken together our nMS results capture a leukotoxin–GPCR complex in solution in a purified system and demonstrate that the F-component HlgB also interacts directly with ACKR1.

**Figure 2.**
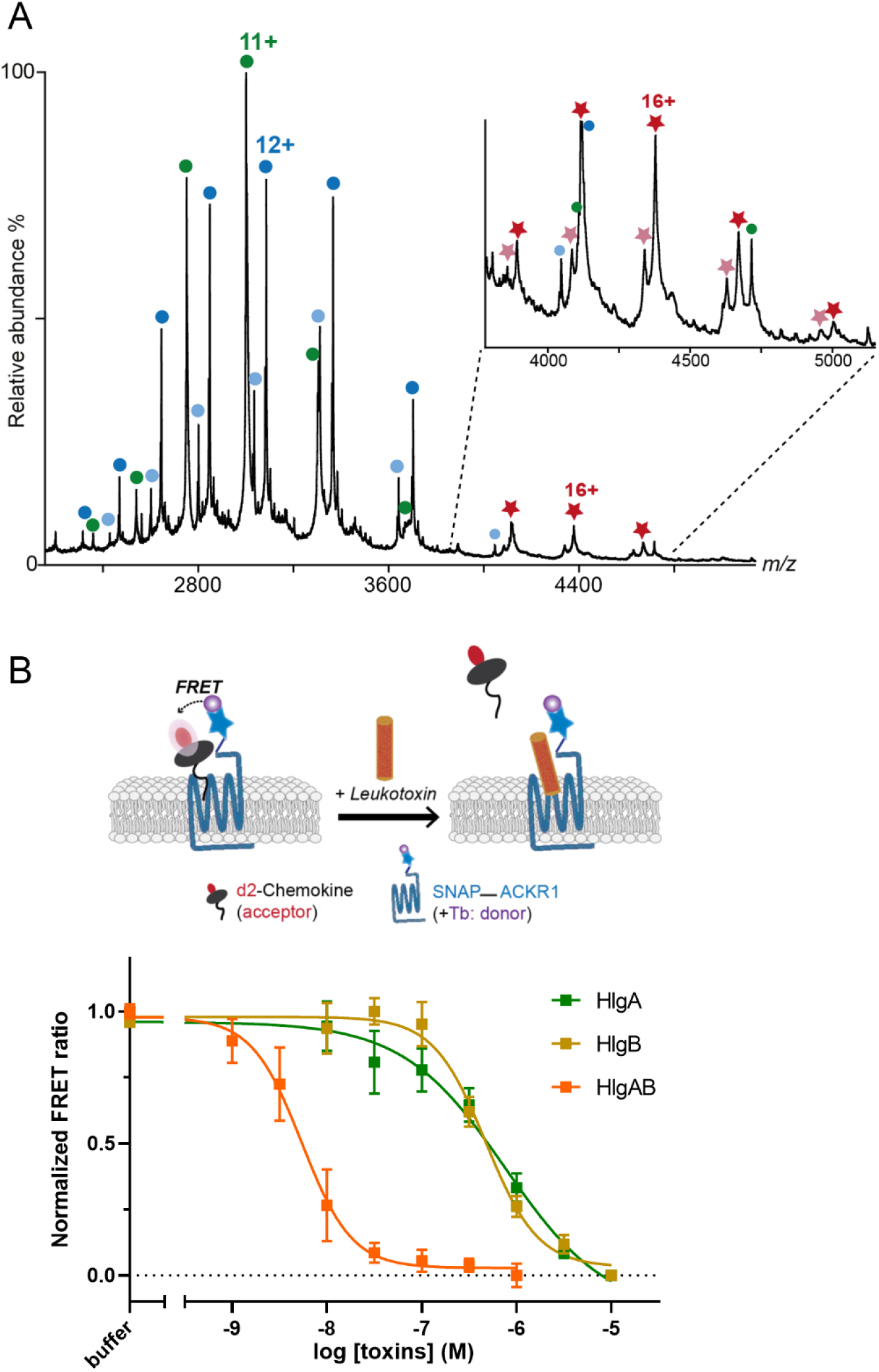
Binding of leukotoxins to ACKR1 *in vitro* and in living cells. A) nMS spectrum of a mixture of 2 μM HlgA and 5 μM ACKR1 treated with PNGase, showing the presence of monomeric HlgA (green circles, 33 004 ± 1 Da), deglycosylated ACKR1 (blue circles, 37 022 ± 1 Da) and partially-hydrolysed deglycosylated ACKR1 (light blue circles, 36 411 ± 1 Da). Complexes formed between HlgA and both forms of deglycosylated ACKR1 are labelled with dark and clear red stars (70 029 ± 2 Da and 69 421 ± 7 Da respectively). B) Scheme explaining TR-FRET competitive binding assay (top) and dose-response curves showing the decrease in TR-FRET ratio between ACKR1 and d2-CCL5 upon addition of HlgA, HlgB and HlgAB equimolar mixture (bottom). Data shown are the mean +/-SEM of one experiment performed in triplicates and are representative of 3 independent experiments. Hill slope values: HlgA -1.1 ± 0.08, HlgB -2.0 ± 0.34, HlgAB -2.1 ± 0.29. IC50 values: HlgA 577 ± 93 nM, HlgB 376 ± 105 nM, HlgAB 6.2 ± 2.7 nM. Values represents the average +/-SD of three independent experiments performed in triplicates.

### Both leukotoxins compete with ACKR1 natural ligand and HlgB binding is cooperative

In order to validate the observed binding *in vitro* and to assess the effect of HlgA and HlgB on ACKR1 binding to an endogenous ligand (CCL5) (Vacchini et al., 2016), we carried out competitive binding assays in living HEK293T cells by Homogenous Time Resolved FRET (TR-FRET) technology (Zwier et al., 2010) (Fig. 2B). SNAP-tag-fused ACKR1 receptor (ST-ACKR1) transiently expressed in HEK293 cells was covalently labelled with Lumi4-terbium as donor. Cells were then incubated in the presence of d2-CCL5, a fluorescent derivative of CCL5 used as ligand tracer. After assay validation (Fig. S4A, B), HlgA and HlgB binding was assessed by competition experiments (Fig 2B). The results revealed that both leukotoxins compete with d2-CCL5 with a slightly higher displacement in the case of HlgB (IC50 = 577 ± 93 and 376 ± 105 nM respectively), confirming the capacity of both toxins to bind to ACKR1 as seen by nMS. Surprisingly, the slopes of the competition curves are consistently different, close to -1 for HlgA and close to -2 for HlgB, suggesting the existence of a positive cooperative binding process with HlgB. This could be correlated with the higher ability of HlgB to homo-dimerize compared to HlgA as observed by nMS with purified toxins. This dimerization was shown to be enhanced in the presence of detergent micelles, and therefore could be more marked at the vicinity of a biological membrane. Finally, binding affinity of an equimolar HlgA and HlgB mixture resulted in a dramatic left shift of the IC50 of the competition curve (6.2 ± 2.7 nM) and the slope of the curve, as observed with HlgB, presented the characteristics of a positive cooperative binding process (Fig. 2B). The observed increased affinity and cooperative binding might originate from an increased avidity, similar to what is observed with bivalent ligands (Vauquelin and Charlton, 2013). Indeed, HlgB is more prone to homo-dimerize compared to HlgA, which may explain the cooperativity observed only for HlgB and the lower IC50 for the latter, whereas a mixture of HlgA and HlgB will lead to hetero-oligomerization with the potential formation of octamers forming the (pre-)pore, resulting in an enhanced affinity through avidity.

### HlgAB and HlgB interfere with ACKR1–ACKR1 interactions in living cells

Based on the observed positive cooperativity for HlgAB and HlgB binding and given that GPCRs are notoriously known to form oligomers including in native tissues (Albizu et al., 2010), we next sought to determine whether the toxins binding could modify receptor oligomerisation. In order to explore these mechanisms, we monitored the effects of leukotoxins on receptor–receptor interactions in living cells using two different resonance energy transfer (RET) strategies: TR-FRET and bioluminescent RET (BRET). BRET is based on the fusion of an energy donor (RLuc or NanoLuc) and an energy acceptor (YFP) to the C-terminus of ACKR1 (Brown et al., 2015) (Fig. 3A), whereas TR-FRET is based on the fusion of a SNAP-tag to ACKR1 extracellular N-terminus that will be labeled with a FRET donor and an acceptor (Maurel et al., 2008) (Fig. 3B). In both techniques, RET signal is sensitive to the distance between the donor and the acceptor while in BRET it can also be sensitive to the relative orientation of the fluorescent probes. The different positioning of the probes can thus sense various receptor rearrangements both in the extracellular and intracellular domains of this GPCR.

**Figure 3.**
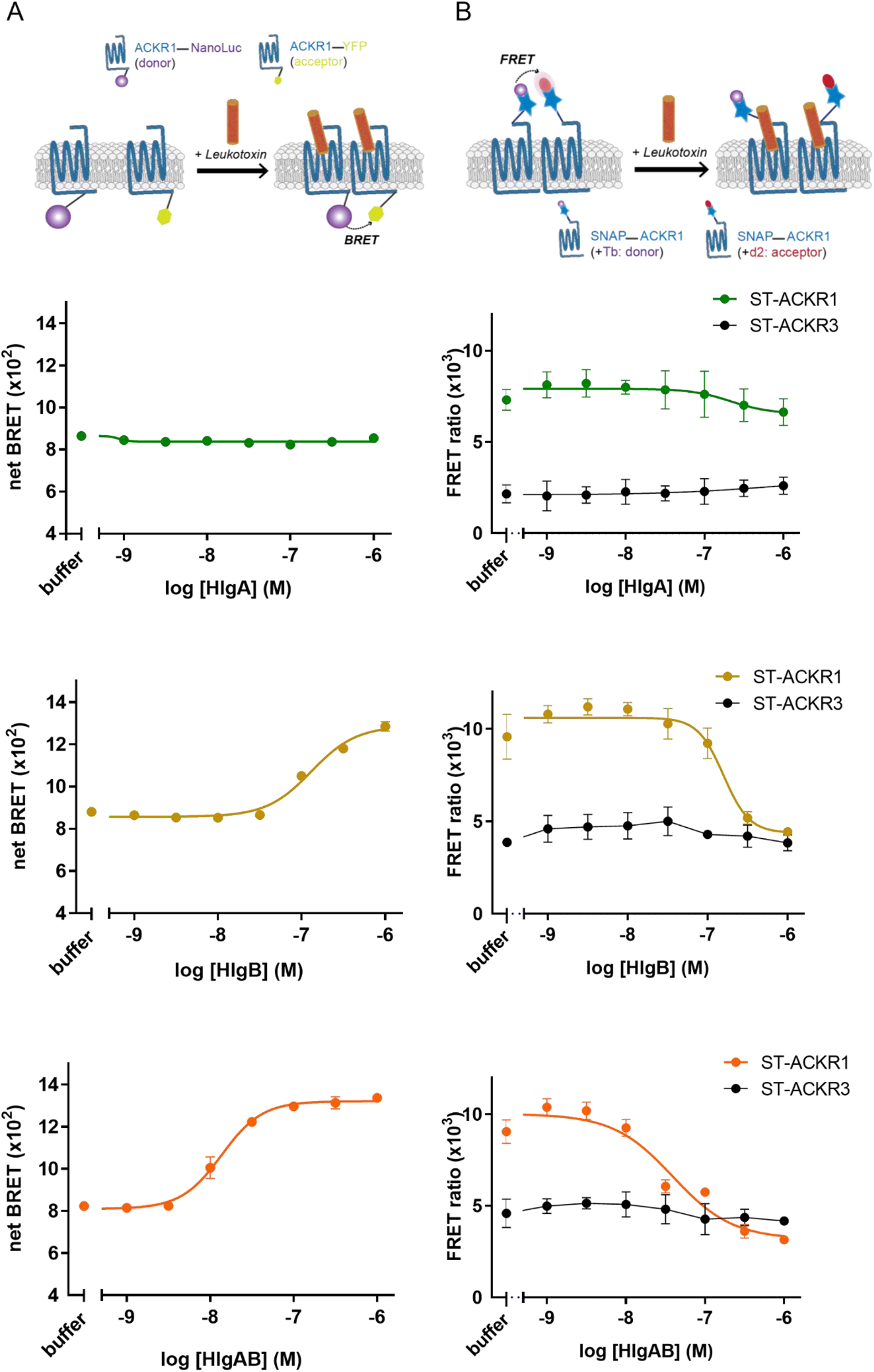
Effect of leukotoxins on ACKR1-ACKR1 interactions in living cells. A) BRET assay showing receptor-receptor interactions at the C-terminal intracellular side of ACKR1. Here also, a constitutive BRET signal is visible prior to adding the different ligands. Adding HlgA (green) does not affect the BRET signal at the tested conditions, whereas HlgB (yellow) and equimolar mixture of HlgAB (orange) induced an increased BRET. Average EC50 of all independent experiments was: 178 ± 28 nM in the presence of HlgB and 18.6 ± 2.5 nM in the presence of HlgAB. B) TR-FRET showing receptor-receptor interactions at the N-terminal extracellular side of ACKR1. A constitutive TR-FRET signal visible prior to adding the different ligands. Adding HlgA (green) does not affect the TR-FRET at the tested conditions, whereas HlgB (yellow) and equimolar mixture of HlgAB (orange) induced a decreased TR-FRET. Average EC50 of all independent experiments was: 429 ± 138 nM in the presence of HlgB and 47 ± 15 nM in the presence of HlgAB. ST, SNAP Tag, ACKR3 was used as control. All data shown are the mean +/-SEM of one experiment performed in triplicates and are representative of at least 3 independent experiments.

We first confirmed the propensity of ACKR1 to form oligomers in living cells. Indeed, saturation of the BRET signal when increasing ACKR1-YFP expression while ACKR1-Nanoluc remained constant strongly supports receptor oligomerisation (Fig. S4C). These results correlate with the presence of purified stable ACKR1 dimers in solution detected by nMS (Fig. 1E) as well as with previous studies confirming the presence of homo-oligomeric ACKR1 by BRET (Chakera et al., 2008). Using the experimental conditions corresponding to the BRET50, we evaluated the impact of HlgA and HlgB binding on BRET signal. HlgA did not induce any modification in the BRET signal while HlgB induced a BRET signal increase (Fig. 3A). Interestingly, HlgAB equimolar mixture resulted in an increased BRET signal, with a tenfold higher potency compared to HlgB (18.6 ± 2.5 nM *versus* 178 ± 28 nM). The one log difference between HlgB and HlgAB is similar to the difference observed in competitive binding experiments (Fig. 2B). The increased BRET signal can be due to an increase in oligomer density and/or to an allosteric conformational rearrangement propagating to the Cter part of ACKR1 leading to a closer proximity and/or a reorientation of the probes, in the presence of HlgAB and HlgB, but not with HlgA at the tested ligand concentrations. Remarkably, this correlates with the conditions showing a cooperative effect in competitive binding assays.

Similar experiments were performed with N-terminus SNAP-tagged receptor that were labelled with Lumi4-Tb (donor) and d2 (acceptor). In the absence of ligand, a constitutive TR-FRET signal was recorded (Fig. 3B), confirming again the capacity ACKR1 to oligomerise. Surprisingly, in contrast to the BRET experiments, the data revealed a decrease in TR-FRET signal between ACKR1 receptors in the presence of HlgB and HlgAB while no significant effect was observed with HlgA at the tested ligand concentrations. Again, the potency of HlgB was 10-fold lower compared to HlgAB (429 ± 138 versus 47 ± 15 nM) in the same range than in the BRET assays. Taken together, the observation of BRET signal increase and TR-FRET signal decrease strongly suggest that HlgB and HlgAB induce large conformational changes leading to a rearrangement of the oligomeric architecture that positions the fluorescent probes further apart in the N-termini and closer together and/or with a different orientation in the C-termini (Fig. 6).

### Conformational changes of ACKR1 upon binding to leukotoxins revealed by HDX-MS

In order to determine the potential conformational changes occurring in ACKR1 monomers upon binding to leukotoxins, we developed an HDX-MS strategy using purified monomeric ACKR1 and leukotoxins *in vitro*. HDX-MS gives information related to solvent accessibility and dynamics of biological complexes. It is based on the exchange kinetics between Deuterium atoms (D) present in the buffer and amide protons (H) of native polypeptide chains in solution (Zheng et al., 2019). Unlike higher resolution approaches, HDX-MS offers the advantage of obtaining dynamic structural information for samples that presents heterogeneous and/or flexible areas. It was successfully applied to characterize the dynamics and interactions of membrane transporters and even GPCRs (Du et al., 2019; Fiorentino et al., 2020; Jia et al., 2020; Möller et al., 2019; Reading et al., 2020; Zhang et al., 2010).

HDX-MS optimization for ACKR1 allowed the identification of 77 peptides covering 70.1 % of WT ACKR1, with 3.41 peptide redundancy (Fig. S5). The highly glycosylated Nter part of ACKR1 was mainly missing, which was compensated when analysing the non-glycosylated N^16,27,33^Q-ACKR1 mutant (Fig. S5). In order to determine the effect of HlgA and HlgB binding on ACKR1 conformational dynamics, we performed differential HDX (ΔHDX) analysis between *apo* ACKR1 and equimolar mixtures of ACKR1: HlgA or HlgB. Biological replicates (2 with WT ACKR1 and 1 with N^16,27,33^Q-ACKR1) were freshly prepared, and deuteration timepoints were performed in triplicates for each condition (Fig. S5). The total numbers of ACKR1 peptides detected in the presence of leukotoxins were lower compared to the receptor analysed alone, most probably due to the overlay with the additional peptides coming from soluble leukotoxins. Mixtures of receptor: leukotoxins were pre-incubated together for 30 min at 4°C prior to HDX-MS analysis.

In order to visualize the leukotoxin-induced changes on ACKR1 and in the absence of a high-resolution structure of the latter, we generated a model for ACKR1 that was validated against HDX-MS data using molecular dynamics simulations (MDS) (Fig. S6, see Methods). Both leukotoxins had similar structural effects on purified monomeric ACKR1, which correlates with the similar binding affinities measured in living cells by TR-FRET. ACKR1 long N-terminal part was mainly protected in the presence of leukotoxins (peptide 9-20 more protected compared to peptide 27-45), whereas HDX data for the majority of ECLs was not present due to the missing sequence coverage (Fig. 4). The overall deuteration of detected TM segments was very low, most likely due to the presence of the detergent micelles surrounding these parts that shield them from the solvent. However, the upper part of H5 (residue 203-215) presented a higher uptake compared to the other TM, and was slightly more protected in the presence of the toxins compared to the receptor alone (Fig. 4). This region also displays higher flexibility in MDS of generated ACKR1 model (Fig. S6). The upper part of TM5 was shown to be important for various CKR–ligand interactions (Arimont et al., 2017), whereas rearrangements in TM5 was shown to play critical roles in signal transmission for various GPCRs (Latorraca et al., 2017). Interestingly, ΔHDX shows that binding of leukotoxins induced allosteric conformational changes that lead to the protection of the H8 and the C-terminal part of ACKR1, as well as both ICL1 and ICL2 (Fig. 4), domains that are known to be critical for Gα binding (Mahoney and Sunahara, 2016). This implies that helix 8 might change its conformation to shield both ICL1 and ICL2 upon binding of ACKR1 to leukotoxins, in a way that blocks G protein accessibility similar to what was observed for Angiotensin 2 receptor AT2R (Zhang et al., 2017). The structural dynamics of H8 is indeed suggested to play an important role in GPCR signalling (Dijkman et al., 2020).

**Figure 4.**
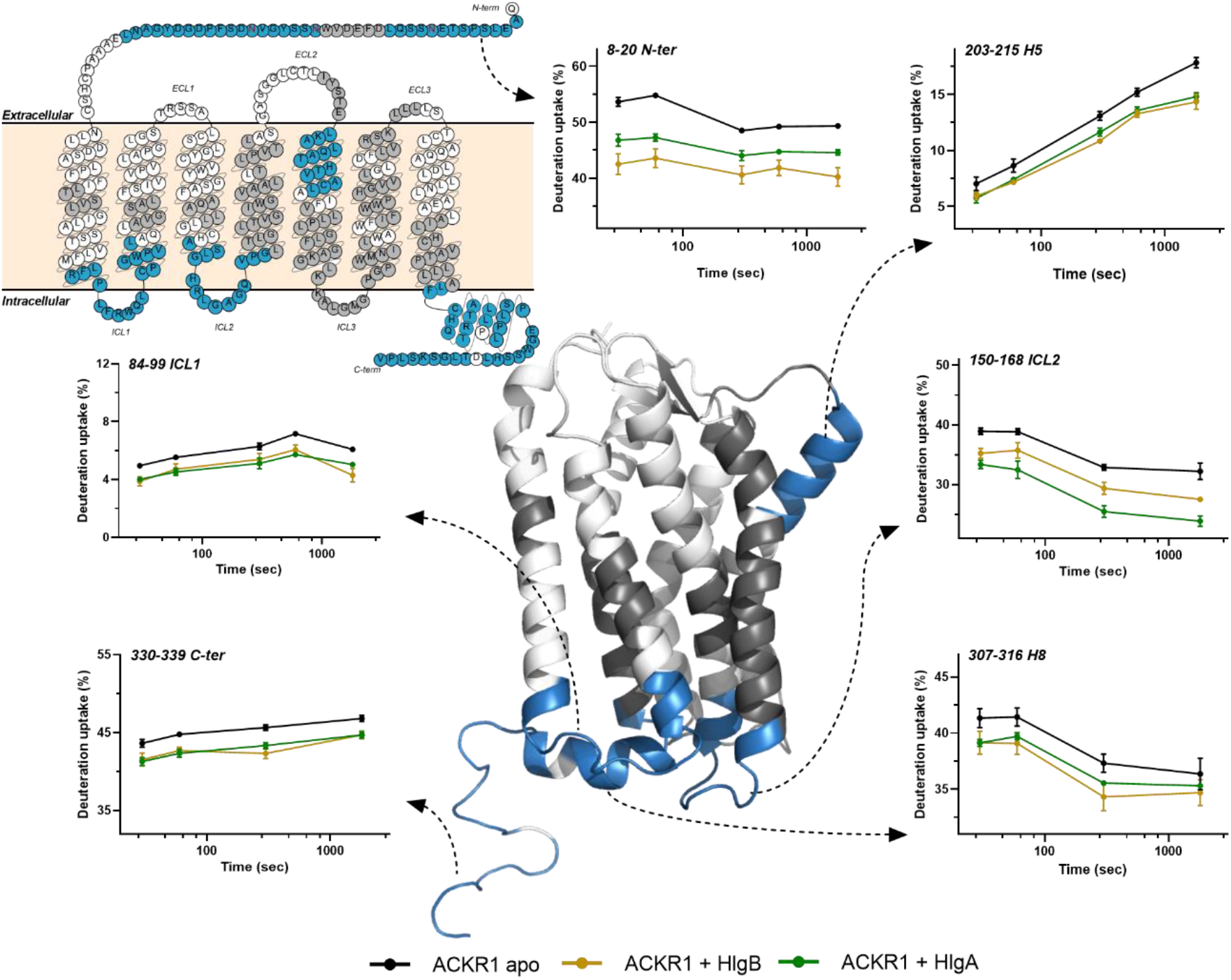
ACKR1 conformational changes upon binding to leukotoxins probed by HDX. HDX results showing statistically significant ΔHDX regions from all three biological replicates, colour-coded onto the snakeplot of ACKR1 adapted from GPCRdb as well as on the structural model generated for ACKR1 where the long N-terminal domain is missing (*c*.*f*. methods). Blue: protected regions, grey: regions with no statistically significant ΔHDX and white: regions with no HDX data. Deuterium uptake of selected peptides is shown for the *apo* receptor (black) and the receptor bound to HlgA (green) or to HlgB (yellow). Uptake plot data are the average and standard deviation for timepoints from n = 3 replicate measurments for one biological preparation of ACKR1.

### HlgB and HlgAB interfere with preassembled ACKR1–Gαi complexes in living cells

The surprising allosteric modulation observed by HDX prompted us to analyse the ACKR1-Gi interactions in living cells. We did not see any G protein activation with ACKR1 in the presence of chemokines CCL2 and CCL5, as demonstrated by the absence of a BRET signal decrease between the α and the βγ subunits of the studied G proteins (Fig. S4D, E). This is not surprising for ACKR1 since it lacks the canonical DRY motif involved in G protein activation (Horuk, 2015). However, when we followed the direct interaction between the C-terminal part of ACKR1 and the Gαi subunit by BRET (Fig. 5A), we observed a constitutive BRET signal in the absence of any ligand, indicating the existence of ACKR1-YFP–Giα1-RLuc complexes in basal conditions (Fig. S4F). This could be similar to the decoy action ACKRs have regarding chemokines present in the extracellular milieu, wherein ACKR1 could also help regulating the concentration of Gα in the intracellular milieu. Adding up to 10 µM of CCL5 or HlgA promoted a weak variation of the BRET signal between ACKR1-YFP and Giα1-RLuc, whereas incubation of cells with increasing concentrations of HlgB or HlgAB strongly reduced basal BRET signal (Fig 5B), suggesting a conformational rearrangement that modifies the distances and/or orientation within the ACKR1–YFP-Giα1-RLuc complexes. Similar to what we observed in all cell-based assays, the effect of HlgAB was nearly 10-fold more potent compared to the effect of HlgB (EC50 796 ± 110 nM and 14 ± 3 nM for HlgB and HlgAB respectively). To further validate the specificity of the observed effect, we performed the same experiments using the mutant Y41F-ACKR1. When mutating the sulfotyrosin Y41, reported as highly important for ACKR1–mediated pore-formation by HlgAB (Spaan et al., 2015), we observed a 10-fold decrease in the effect of HlgAB (Fig. 5C). Interestingly, mutating Y30, another potentially sulfated tyrosine present at the Nter, did not result in any significant effect on ACKR1-Gαi1 binding, implying that this site is either not sulfated or that sulfation of this tyrosine is not important for binding to leukotoxins, as reported previously (Spaan et al., 2015). These results thus demonstrate that ACKR1–Giα1 architecture can be modified in living cells upon leukotoxins binding, confirming the biological relevance of the allosteric modifications observed by HDX-MS.

**Figure 5.**
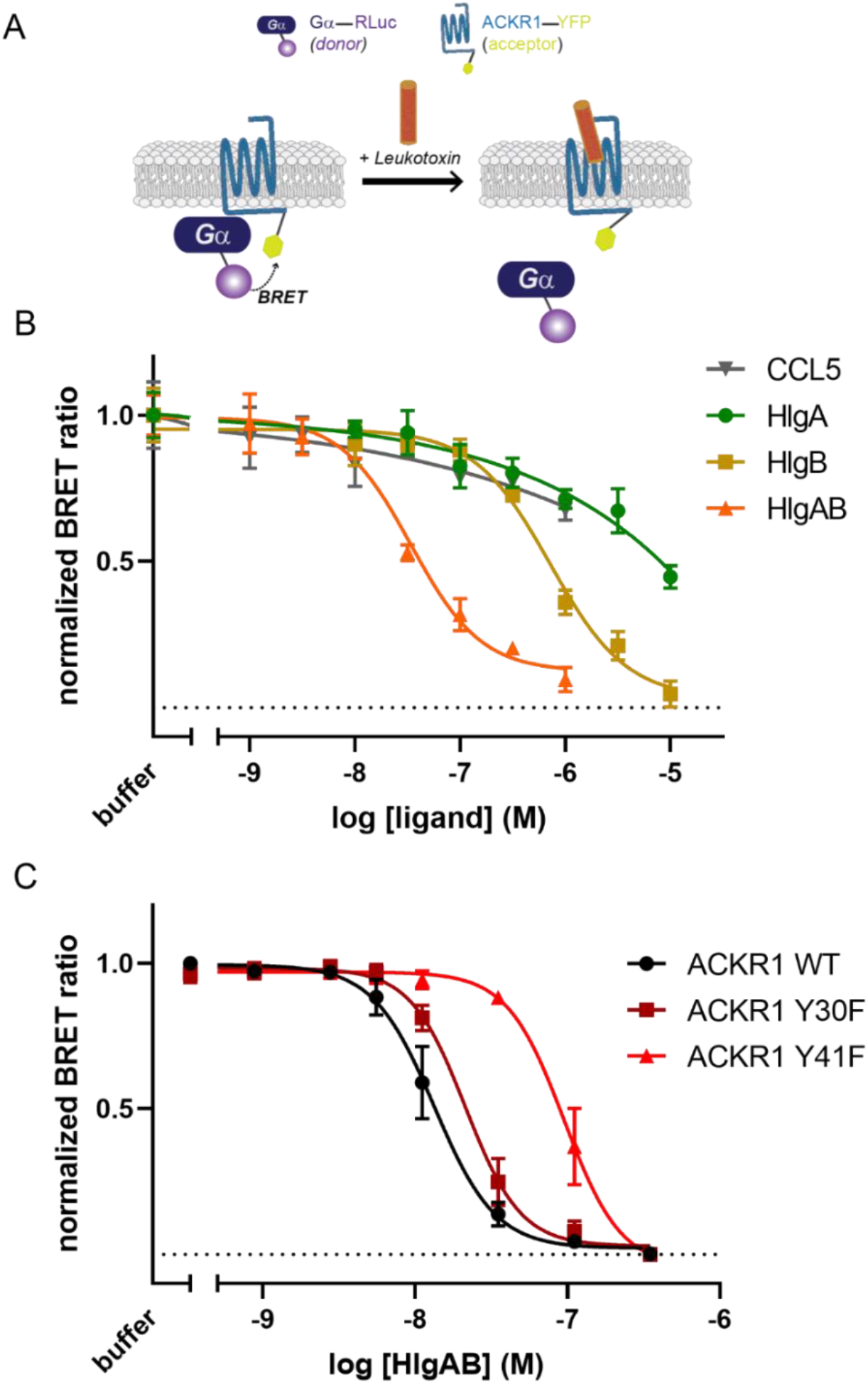
HlgB and HlgAB interfere with preassembled ACKR1-Gαi complexes in living cells. A) Scheme explaining BRET assay used to probe ACKR1-Gα interactions. B) Normalised BRET assay between ACKR1–YFP and Gαi1-RLuc showing that CCL5 and HlgA has no specific effect on ACKR1-Gαi1 interactions at the tested concentrations, whereas adding HlgB or HlgAB resulted in the dissociation of ACKR1-Gαi1 complexes in living cells. EC50 was 796 ± 110 nM for HlgB and 13.5 ± 3.2 nM for HlgAB. C) Normalised BRET assay showing the effect of HlgAB mixture on ACKR1–Gαi1 interactions for WT, Y30F and Y41F ACKR1. Effect of HlgAB on the dissociation between ACKR1 and Gαi1 decreased when mutanting Y41, evidenced by an EC50 increase up to 106 ± 42 nM, whereas the EC50 did not change significantly when mutating Y30 (21.7 ± 5.3 nM). Cell-based assays data shown are the mean +/-SEM of one experiment performed in triplicates and are representative of 3 independent experiments.

## CONCLUSION

SA leukotoxins targeting GPCRs represents an attractive aspect in modulating GPCR function and remains largely unexplored. We chose to focus on ACKR1 since it is a crucial target for SA pathogenesis, being not only expressed in myeloid cells like the other targeted receptors, but also in erythrocytes and endothelial cells. We demonstrate that HlgB also recognises ACKR1, making it the second F-component leukotoxin with an identified receptor. Our results may explain the observation that HlgB binds to erythrocytes independently from HlgA (Ozawa et al., 1995). Both leukotoxins were able to compete with ACKR1 natural ligand, however with a difference in the mechanism. The cooperativity that accompanies toxin ability to dimerize and oligomerize seems a key factor that drives the effects observed on ACKR1 conformational changes. While HlgB shows a higher ability to homodimerize compared to HlgA, the presence of both HlgA and HlgB at the vicinity of a membrane expressing their targeted receptor induces the formation of an octameric pore (Yamashita et al., 2014, 2011). Dimerisation and oligomerisation increases the avidity in the system which in turns increases the effects of HlgBB and (HlgAB)_4_ on ACKR1.

The proposed model describing the first steps of pore-formation by HlgAB should thus be reconsidered. We propose that both HlgA and HlgB recognise ACKR1, and their binding to the receptor will induce conformational changes at the extracellular N-terminal part as well as allosteric changes at the C-terminal part in H8 that shields G protein binding sites in the ICLs (Fig. 6). Leukotoxin binding will therefore dissociate ACKR1-G protein complexes, releasing available G protein in the intracellular milieu. The capacity of HlgB and HlgAB to dimerize will lead to changes in ACKR1-ACKR1 constitutive interactions in living cells. However, only the presence of HlgAB heterodimer can lead to pore-formation. Further studies will thus be important to shed the light on the mechanisms behind conformational changes of both leukotoxins and the potential role of the GPCR receptor during the different pore-formation steps.

**Figure 6.**
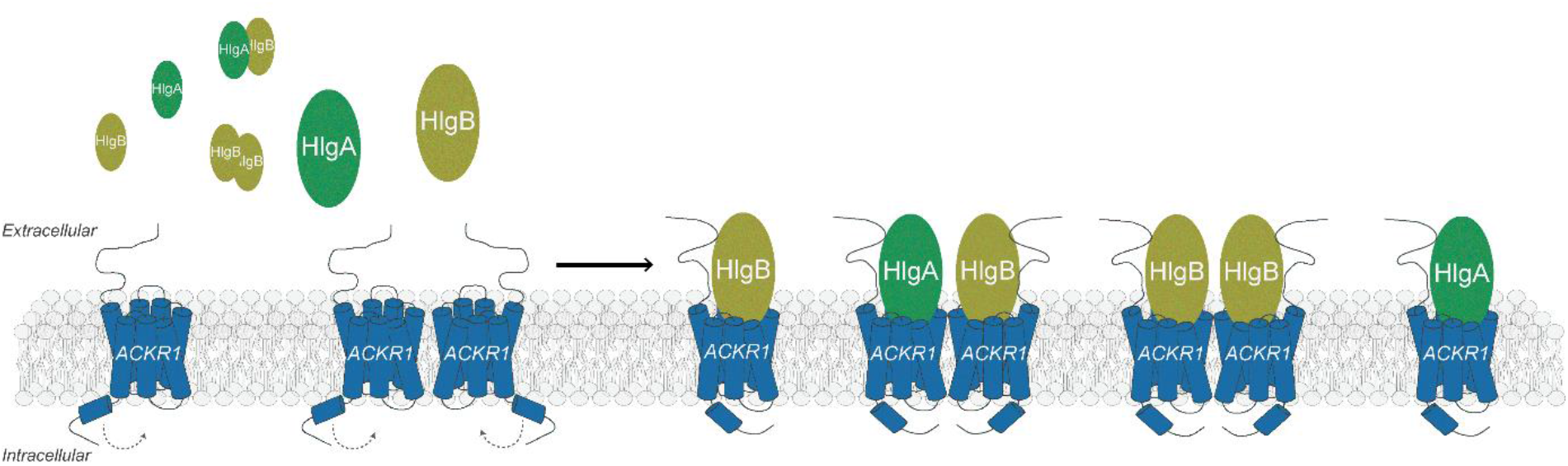
Proposed first steps of pore-formation by HlgAB. ACKR1 (blue) is present in both monomeric and dimeric forms in cellular membranes. Soluble HlgA (green) and HlgB (olive) secreted by SA can be present in monomeric and dimeric forms. Both leukotoxins recognise the cellular membrane by specific interactions with ACKR1, and interaction of each toxin with ACKR1 will lead to conformational changes at both N-and C-termini of the GPCR. HlgB homodimers and HlgAB heterodimers could interfere with receptor-receptor interactions, but only HlgAB-(ACKR1)_2_ complex will lead to the formation of the pore.

## ACKNOWLEDGMENTS

The authors would like to thank all members of the Granier-Mouillac team at the IGF for fruitful discussions, and Isabel Brabet for her help with the cell-based assays. CB would like to thank James Sturgis and all the members of the “Plateforme Protéomique” at the IMM in Marseille for giving her access to the Synapt for preliminary data acquisitions. The authors are grateful for the regional funds (FEDER, Région Occitanie) and funds from MUSE and the Labex EpiGenMed for the Synapt G2-Si and the automated HDX platform. Mass spectrometry experiments were carried out using the facilities of the Montpellier Proteomics Platform (PPM, BioCampus Montpellier) and the Plateforme Protéomique (IMM, Marseille). All cell-based assays were carried out at the Arpège facility (BioCampus, Montpellier). This work was supported by the french Agence Nationale de la Recherche (project ANR-17-CE15-0002-01, CHEMSPEC).

## AUTHOR CONTRIBUTIONS

CG, SG and CB designed the research. CG, CL, PL, SJ, SF, FP and RS designed, expressed and purified the proteins. CG and JH performed and analysed cell-based assays with the help of TD. EDN performed and analysed HDX data with CB. CL performed modelling and computational analysis of ACKR1. CB performed and analysed native MS experiments. SG and CB supervised research, analysed data and CB wrote the manuscript with the help of SG and contributions from all authors.

## METHOD DETAILS

### Protein constructs

For production in insect or human cells, the full-length synthetic gene of human ACKR1 isoform-2 (UniprotKB-Q16570) was subcloned into modified pFastBac or pCMV-Dest vectors (Thermofischer) respectively, resulting in the full-length ACKR1 bearing the influenza virus hemagglutinin (HA) signal peptide followed by a Flag epitope at N-terminal. For human cell-based assays, the full-length synthetic gene of human ACKR1 was subcloned into pcDNA3.1-YFP vector in frame with the N-terminus of YFP resulting in ACKR1-YFP, into pcDNA3.1-Nanoluc vector in frame with the N-terminus of Nanoluc resulting in ACKR1-Nanoluc, and into pcDNA3.1-SNAP vector in frame with the C-terminus of SNAP tag resulting in SNAP-ACKR1. The synthetic genes of mature *Staphylococcus aureus* HlgA (UniprotKB-P0A074) residue 30-309 and HlgB (UniprotKB-P0A077) residue 25-325 were subcloned into popinE vectors (OPPF-UK) resulting in HlgA and HlgB bearing a C-terminal (His)_6_-tag.

### Expression and purification of ACKR1

Expression of ACKR1 was carried out in insect and human cells. ACKR1 constructs were expressed in HEK293 GnTI^-^cells (ATCC) using BacMam baculovirus transduction. HEK cells were grown in suspension in Ex-Cell^®^ 293 Serum-Free Medium (Sigma) with 2% FBS and were infected at a density of 2×10^6^ cells/mL using a 1/10 (v/v) baculovirus solution. Culture flasks were shaken for 72 h at 37°C with 5% CO_2_ and a solution of 5 mM sodium butyrate was added to the culture flasks 24 h post-infection. Cells were harvested by centrifugation (3,000 rpm) 72 h post-infection and cell pellets were stored at -80 °C until purification. ACKR1 constructs were also expressed in Sf9 insect cells (Life Technologies) using the pFastBac baculovirus system (Thermofischer). Sf9 cells were grown in Ex-Cell^®^ 420 Medium (Sigma) and were infected at a density of 4×10^6^ cells/mL using a 1/200 (v/v) baculovirus solution. Culture flasks were shaken for 48 h at 28 °C and cells were harvested by centrifugation (3,000 rpm), and cell pellets were stored at – 80 °C until purification. Purification was carried out in similar conditions regardless cell expression. After thawing the frozen cell pellets, cells were lysed by osmotic shock adding 1/10 (v/v) lysis buffer consisting of 10 mM Tris (pH 7.5), 1 mM EDTA and containing 2 mg/mL of iodoacetamide (Sigma) and protease inhibitors (Leupeptin (Euromedex), Benzamidine and PMSF (Sigma)). Lysed cells were centrifuged (16,000 rpm) and the membrane pellets were suspended in a 1/20 (v/v) solubilisation buffer consisting of 50 mM HEPES (pH 7.5), 150 mM NaCl, 0.5% (w/v) n-dodecyl-D-maltoside (DDM, Anatrace), 0.1% (w/v) cholesteryl-hemi-succinate (CHS, Sigma), 2 mg/mL of iodoacetamide and protease inhibitors. Receptors were extracted from the membrane pellets using a glass dounce grinder and the extracted mixture was stirred for 1 h at 4 °C, then centrifuged (16,000 rpm). The supernatant was loaded by gravity flow onto anti-Flag M2 antibody resin. The resin was washed with 10 column volumes (CV) of a DDM wash buffer consisting of 50 mM HEPES (pH 7.5), 150 mM NaCl, 0.1% (w/v) DDM and 0.02% (w/v) CHS. For native MS experiments, DDM concentration was decreased to reach 2 critical micelle concentration (CMC) using 10 CV of wash buffer 2 made up of 50 mM HEPES (pH 7.5), 150 mM NaCl, 0.025% (w/v) DDM and 0.005% (w/v) CHS. The bound receptor was eluted in the wash buffer 2 supplemented with 0.4 mg/mL Flag-peptide. For HDX-MS analysis, detergent was changed from DDM to lauryl maltose neopentyl glycol (LMNG) using LMNG exchange buffer containing 50 mM HEPES (pH 7.5), 150 mM NaCl, 0.2% (w/v) LMNG and 0.01% (w/v) CHS. The detergent exchange was performed by washing the column with a series of 5 buffers (3 CV each) made up of the following ratios (v/v) of LMNG exchange buffer and DDM wash buffer: 1/3, 1/1, 3/1, 9/1, 1/0. Additional 10 CV wash was performed to decrease the detergent concentration to 2CMC LMNG using 50 mM HEPES (pH 7.5), 150 mM NaCl, 0.02% (w/v) LMNG and 0.001% (w/v) CHS followed by the last wash LMNG buffer consisting of 50 mM HEPES (pH 7.5), 150 mM NaCl, 0.002% (w/v) LMNG and 0.0001% (w/v) CHS. The bound receptor was eluted in the last wash LMNG buffer supplemented with 0.4 mg/mL Flag-peptide. The eluted solution of receptors was concentrated to 500 μL using 50 kDa spin filters (Millipore) and further purified by size exclusion chromatography on a Superdex 200 Increase 10/300 column (GE Healthcare) in the last wash buffer. For PNGase F (NEB) treatment, 5 µL of enzyme at 500 000 U/mL were added to 0.2 mg of ACKR1 and incubated overnight at 4°C. Receptor was purified by size exclusion chromatography as mentioned previously. Fractions containing monodisperse ACKR1 were collected and directly analysed for native MS or pooled and concentrated for HDX analyses.

### Expression and purification of HlgA and HlgB

Toxins were expressed with (His)_6_-tags at their C-termini in competent C43 (DE3) *Escherichia coli* cells (NEB) for HlgA and in BL21 (DE3) *Escherichia coli* cells (NEB) for HlgB. Transformed cells were grown at 37°C in Terrific broth for HlgA and in LB broth for HlgB supplemented with 100 µg/mL ampicillin to a density of OD600 = 0.6. Expression was then induced for 6 hrs at 22°C by addition of 0.5 mM IPTG. Cells were harvested by centrifugation (3,000 rpm) and cell pellets were stored at – 80°C until purification. After thawing the frozen cell pellets, cells were lysed by sonication in a lysis buffer consisting of 20 mM Tris (pH 8), 300 mM NaCl, 2 mg/mL of iodoacetamide (Sigma) and protease inhibitors (Leupeptin (Euromedex), Benzamidine and PMSF (Sigma)). Lysed cells were centrifuged (16,000 rpm) and the supernatant was adjusted to 40 mM imidazole and loaded onto a nickel NTA Agarose resin. The resin was washed with 10 CV of wash buffer 1 consisting of 50 mM HEPES (pH 7.5) and 1 M NaCl, and with 10 CV of wash buffer 2 consisting of 50 mM HEPES (pH 7.5) and 150 mM NaCl supplemented with 40 mM imidazole. Bound (His)_6_-toxins were eluted with wash buffer 2 supplemented with 200 mM imidazole. The eluted solution of toxins was concentrated to 500 μL using 30 kDa spin filters (Millipore) and further purified by size exclusion chromatography on a Superdex 200 Increase 10/300 column (GE Healthcare) in 50 mM HEPES (pH 7.5) and 150 mM NaCl. Fractions containing monodisperse toxins were collected and concentrated for native MS and HDX analyses and cell-based assays.

### Native mass spectrometry

Prior to MS analysis, proteins were buffer exchanged into 200 mM ammonium acetate buffer pH 7.4 (Sigma), supplemented with 0.02% DDM for ACKR1 and for toxins: detergent interactions analysis, using Bio-Spin microcentrifuge columns (Bio-Rad Laboratories). Intact MS spectra were recorded on a Synapt G2-Si HDMS instrument (Waters Corporation) modified for high mass analysis and operated in ToF mode. Samples were introduced into the ion source using borosilicate emitters (Thermo Scientific). Optimized instrument parameters for ACKR1 alone or in the presence of the toxins were as follows: capillary voltage 1.4 kV, sampling cone voltage 150 V, offset voltage 80 V, transfer collision voltage 25 V, argon flow rate 8 mL/min and trap bias 25 V. Collision voltage in the trap was optimized between 50 and 110 V depending on the sample, and to the minimum activation required to strip the detergent micelle when it was present. Data was processed using MassLynx v.4.2 (Waters) and UniDec (Marty et al., 2015).

### Transfection for cell-based assays

HEK293 cells (ATCC) were grown in Dulbecco’s Modified Eagle Medium supplemented with 10% FBS and 1% Penicillin/streptomycin at 37°C with 5% CO_2_. Transfection was performed using lipofectamine 2000 (Life technology) in polyornithine coated black-walled, dark-bottom 96-well plates. The quantity of transfected DNA was optimised for each construct and 150 ng of DNA in total, supplemented by empty PRK5, were added to 50.10^4^ cells/well for transfection.

### Competitive binding assay by TR-FRET

1.5 ng of SNAP-ACKR1 were added to 50.10^4^ cells/well for transfection. 48 h post-transfection, the plate was washed twice with TagLite^®^ (Cisbio, Codolet, France) and the extracellular SNAP-receptors were labelled with SNAP-Lumi4-Tb (100 nM, Cisbio, Codolet, France) for 1 h at 37°C. The plate was washed three times with TagLite^®^. 12 nM of d2-labelled CCL5 and increasing concentration (0, 1, 3.6, 10, 31.6, 100, 316, 1000, 3160 and 10000 nM) of toxins, HlgA, HlgB or an equimolar mixture of HlgA and HlgB, were added to cells. Lumi-4-fluorescence signals were osberved on a Pherastar plate reader (BMG Labtech): samples were illuminated at 337 nm and fluorescence was acquired at 620 nm (donor) and 665 nm (TR-FRET) over time. The ratio of the signals (665/620) was plotted versus the toxin concentration. Dose-response curves were generated using GraphPad Prism 6^®^ (GraphPad Software, Inc., San Diego, CA).

### Receptor-receptor interactions by BRET

80 ng of ACKR1-YFP and 10 ng of ACKR1-NanoLuc were added to 50.10^4^ cells/well for transfection. 24 h post-transfection, the plate was washed three times with Phosphate Buffered Saline (PBS, Sigma) and then cells are pre-incubated 5 minutes at 37°C with 5 μM coelenterazine h (Promega) before adding ligands diluted in PBS supplemented with 0.9 mM CaCl_2_ and 0.5 mM MgCl_2_. Increasing concentration (0, 1, 3.6, 10, 31.6, 100, 316, 1000, 3160 and 10000 nM) of toxins, HlgA, HlgB or an equimolar mixture of HlgA and HlgB, were added to cells. BRET readings were collected using a Mithras 2 plate reader (Berthold Technologies GmbH, Bad Wildbad, Germany) with 0.1 seconds integration time per well. The reading chamber was maintained at 37°C throughout the entire reading time. The BRET signal was calculated by the ratio of emission of YFP (535nm) to NanoLuc (460nm). Dose-response curves were generated using GraphPad Prism 6^®^ (GraphPad Software, Inc., San Diego, CA).

### Receptor-receptor interactions by TR-FRET

50 ng of SNAP-ACKR1 or SNAP-ACKR3 (control) were added to 50.10^4^ cells/well for transfection. 24 h post-transfection, the plate was washed twice with TagLite^®^ (Cisbio, Codolet, France) and the extracellular SNAP-receptors were labelled with SNAP-Lumi4-Tb (100 nM, Cisbio, Codolet, France) and SNAP-red (300 nM, Cisbio, Codolet, France) for 1 h at 37°C. The plate was washed four times with PBS. Increasing concentration (0, 1, 3.6, 10, 31.6, 100, 316, 1000, 3160 and 10000 nM) of toxins, HlgA, HlgB or an equimolar mixture of HlgA and HlgB, were then added to cells. 30 minutes after addition of leukotoxins, TRF readings were collected using a SPARK 20M plate reader (TECAN) with 150 µs of lag time and 500 µs of integration time per well. Excitation at 340 nm and emission at 620 nm (donor) and 665 nm (TR-FRET) over time. The ratio of the signals (665/620) was plotted versus the concentrations of toxin.

### Receptor-Gα interactions by BRET

To follow interaction between ACKR1 and Gα subunit, 40 ng of ACKR1-YFP and 10 ng of Gα-RLuc were added to 50.10^4^ cells/well for transfection. To follow G protein activation, cells were transfected with each ST-Receptor (10 ng CCR5, 10 ng CCR2, 20 ng ACKR1) and in parallel with 10 ng Gαi1-RLuc, 10 ng β2 and 20 ng γ-Venus. 24 h post-transfection, the plate was washed three times with Phosphate Buffered Saline (PBS, Sigma) and then cells were pre-incubated 5 minutes at 37°C with 5 μM coelenterazine h (Promega) before adding ligands diluted in PBS supplemented with 0.9 mM CaCl_2_ and 0.5 mM MgCl_2_. BRET readings were collected using a Mithras 2 plate reader (Berthold Technologies GmbH, Bad Wildbad, Germany) with 0.1 seconds integration time per well. The reading chamber was maintained at 37°C throughout the entire reading time. The BRET signal was calculated by the ratio of emission of YFP (535 nm) to RLuc (480 nm). Dose-response curves were generated using GraphPad Prism 6^®^ (GraphPad Software, Inc., San Diego, CA).

### Structural modeling of ACKR1

Starting models of ACKR1 were obtained by submitting residues 51-326 of ACKR1 sequence to the GPCR-I-TASSER (Zhang et al., 2015) and to the GPCRM Structure modeling webservers (Miszta et al., 2018). The best model generated by each approach was then subjected to molecular dynamics simulations (MDS), in order to refine them and assess their stability, dynamics and correlation with experimental HD/X profiles. The starting I-TASSER model displayed local unfolding at the top of TM1 and at the bottom of TM6 and the backbone geometry of these regions was manually idealized in coot (Emsley et al., 2010). The GPCR-I-TASSER and GPCRM MD systems were then set up using the CHARMM-GUI membrane builder (Lee et al., 2016). The starting models were each inserted into a hydrated, equilibrated bilayer composed of 163 molecules of 2-Oleoyl-1-palmitoyl-sn-glycero-3-phosphocholine and 20 molecules of cholesterol. Na^+^ and Cl^-^ions were added to neutralize the system and reach a final concentration of 150 mM. MD calculations were performed in GROMACS2020 (Pronk et al., 2013) using the CHARMM36m force field (Huang et al., 2017) and the CHARMM TIP3P water model. The input systems were subjected to energy minimization, equilibration and production MDS using the CHARMM-GUI input scripts for GROMACS. For both starting models, we performed two production runs with or without secondary structure (SS) restraints. The aim of these SS restraints was to enable potential tertiary structure re-arrangements without disrupting existing SS elements. For the GPCRM model, we generated distance restraints between amide O and HN atoms that were within 2.3 Å of each other in the starting structure, while excluding the loop regions. Production MDS were then run for 1.3 and 0.6 µs for the unrestrained and restrained systems, respectively. For the GPCR-I-TASSER model, we similarly applied SS restraints within the TMs, but also within the ECL2 β hairpin which was formed in the starting model. In addition, the SS restraints were generated using a model pre-equilibrated through 80 ns of unrestrained MDS, which was then used as the starting point for the restrained MDS. This was done to allow the GPCR-I-TASSER model to fully relax after geometry optimization of TM1 and 6. Production MDS were then run for 1.8 and 1.3 µs for the unrestrained and restrained systems, respectively. Subsequently the trajectories were analyzed using GROMACS tools to yield root mean square deviations (RMSD) and root mean square fluctuations (RMSF). In order to correlate MDS with HDX-MS data, we calculated for each trajectory the average main chain solvent accessible surface area (SASA) and H-bonds corresponding to the HDX peptides (with the exception of peptides 34-45, 330-339, 332-339 and 333-339 located in the intrinsically disordered termini). The RMSF, SASA and H-bonds profiles from all MDS were mostly consistent with the observed relative deuterium uptake, however qualitative agreement was best for the unrestrained I-TASSER model, in particular in the TM5 and H8 regions (Fig. S6). Consequently, a representative snapshot from this trajectory was selected for mapping of ΔHDX data (Fig. 4).

### Hydrogen-Deuterium exchange mass spectrometry

HDX-MS experiments were performed using a Synapt G2-Si HDMS coupled to nanoAQUITY UPLC with HDX Automation technology (Waters Corporation). ACKR1 in LMNG detergent was concentrated up to 10-20 µM and optimization of the sequence coverage was performed on undeuterated controls. Analysis of freshly prepared ACKR1 apo, ACKR1: HlgA and ACKR1: HlgB (1: 1 ratio) mixtures were performed as follows: 3 µL of sample are diluted in 57 µL of undeuterated for the reference or deuterated last wash SEC buffer. The final percentage of deuterium in the deuterated buffer was 95%. Deuteration was performed at 20°C for 30, 60, 300, 600 and 1800 sec. Next, 50 µL of reaction sample are quenched in 50 µL of quench buffer (KH_2_PO_4_ 50 mM and K_2_HPO_4_ 50mM, pH 2.3) at 0°C. 80 µL of quenched sample are loaded onto a 50 µL loop and injected on an Enzymate pepsin column (300 Å, 5 µm, 2.1 mm X 30 mm, Waters) maintained at 15°C, with 0.2% formic acid at a flowrate of 100 µL/min and an additional backing pressure of 6000 psi controlled by the HDX regulator kit (Waters). The peptides are then trapped at 0°C on a Vanguard column (ACQUITY UPLC BEH C18 VanGuard Pre-column, 130Å, 1.7 µm, 2.1 mm X 5 mm, Waters) for 3 min, before being loaded at 40 µL/min onto an Acquity UPLC column (ACQUITY UPLC BEH C18 Column, 1.7 µm, 1 mm X 100 mm, Waters) kept at 0°C. Peptides are subsequently eluted with a linear gradient (0.2% formic acid in acetonitrile solvent at 5% up to 35% during the first 6 min, then up to 40% and 95% over 1 min each) and ionized directly by electrospray on a Synapt G2-Si mass spectrometer (Waters). HDMS^E^ data were obtained by 20-30 V trap collision energy ramp. Lock mass accuracy correction was made using a mixture of leucine enkephalin. The pepsin column was then washed three times with Guanidine-HCl 1.5 M, acetonitrile 4% and formic acid 0.8% and a blank is performed between each sample in order to minimize the carry-over. All timepoints were performed in triplicates.

Peptide identification was performed from undeuterated data using ProteinLynx global Server (PLGS, version 3.0.3, Waters). Peptides are filtered by DynamX (version 3.0, Waters) using the following parameters: minimum intensity of 1000, minimum product per amino acid of 0.2, maximum error for threshold of 5 ppm. All peptides were manually checked and data was curated using DynamX. Back exchange was not corrected since we are measuring differential HDX and not absolute one. Statistical analysis of all ΔHDX data was performed using Deuteros 2.0 (Lau et al., 2020) and only peptides with a 99% confidence interval were considered. MS^E^ raw data will be deposited to the ProteomeXchange Consortium via PRIDE.

## REFERENCES

Albizu, L., Cottet, M., Kralikova, M., Stoev, S., Seyer, R., Brabet, I., Roux, T., Bazin, H., Bourrier, E., Lamarque, L., et al. (2010). Time-resolved FRET between GPCR ligands reveals oligomers in native tissues. Nat. Chem. Biol. 6, 587–594.

Allison, T.M., and Bechara, C. (2019). Structural mass spectrometry comes of age: new insight into protein structure, function and interactions. Biochem. Soc. Trans. 47, 317–327.

Alonzo, F., and Torres, V.J. (2014). The Bicomponent Pore-Forming Leucocidins of Staphylococcus aureus. Microbiol. Mol. Biol. Rev. 78, 199–230.

Alonzo III, F., Kozhaya, L., Rawlings, S.A., Reyes-Robles, T., DuMont, A.L., Myszka, D.G., Landau, N.R., Unutmaz, D., and Torres, V.J. (2012). CCR5 is a receptor for Staphylococcus aureus leukotoxin ED. Nature 493, 51–55.

Arimont, M., Sun, S.-L., Leurs, R., Smit, M., de Esch, I.J.P., and de Graaf, C. (2017). Structural Analysis of Chemokine Receptor–Ligand Interactions. J. Med. Chem. 60, 4735–4779.

Assis, L.M., Nedeljković, M., and Dessen, A. (2017). New strategies for targeting and treatment of multi-drug resistant Staphylococcus aureus. Drug Resist. Updat. 31, 1–14.

Bechara, C., and Robinson, C.V. (2015). Different Modes of Lipid Binding to Membrane Proteins Probed by Mass Spectrometry. J. Am. Chem. Soc. 137, 5240–5247.

Brown, N.E., Blumer, J.B., and Hepler, J.R. (2015). Bioluminescence Resonance Energy Transfer to Detect Protein-Protein Interactions in Live Cells. In Protein-Protein Interactions, C.L. Meyerkord, and H. Fu, eds. (New York, NY: Springer New York), pp. 457–465.

Calabrese, A.N., and Radford, S.E. (2018). Mass spectrometry-enabled structural biology of membrane proteins. Methods 147, 187–205.

Cancellieri, C., Vacchini, A., Locati, M., Bonecchi, R., and Borroni, E.M. (2013). Atypical chemokine receptors: from silence to sound. Biochem. Soc. Trans. 41, 231–236.

Chakera, A., Seeber, R.M., John, A.E., Eidne, K.A., and Greaves, D.R. (2008). The Duffy Antigen/Receptor for Chemokines Exists in an Oligomeric Form in Living Cells and Functionally Antagonizes CCR5 Signaling through Hetero-Oligomerization. Mol. Pharmacol. 73, 1362–1370.

Dijkman, P.M., Muñoz-García, J.C., Lavington, S.R., Kumagai, P.S., dos Reis, R.I., Yin, D., Stansfeld, P.J., Costa-Filho, A.J., and Watts, A. (2020). Conformational dynamics of a G protein–coupled receptor helix 8 in lipid membranes. Sci. Adv. 6, eaav8207.

Du, Y., Duc, N.M., Rasmussen, S.G.F., Hilger, D., Kubiak, X., Wang, L., Bohon, J., Kim, H.R., Wegrecki, M., Asuru, A., et al. (2019). Assembly of a GPCR-G Protein Complex. Cell 177, 1232-1242.e11.

Emsley, P., Lohkamp, B., Scott, W.G., and Cowtan, K. (2010). Features and development of Coot. Acta Crystallogr. D Biol. Crystallogr. 66, 486–501.

Engen, J.R., Botzanowski, T., Peterle, D., Georgescauld, F., and Wales, T.E. (2020). Developments in Hydrogen/Deuterium Exchange Mass Spectrometry. Anal. Chem. 93, 567–582.

Fiorentino, F., Sauer, J.B., Qiu, X., Corey, R.A., Cassidy, C.K., Mynors-Wallis, B., Mehmood, S., Bolla, J.R., Stansfeld, P.J., and Robinson, C.V. (2020). Dynamics of an LPS translocon induced by substrate and an antimicrobial peptide. Nat. Chem. Biol. 17, 187–195.

Foster, T.J. (2004). The Staphylococcus aureus “superbug.” J. Clin. Invest. 114, 1693–1696.

Hansell, C.A.H., Hurson, C.E., and Nibbs, R.J.B. (2011). DARC and D6: silent partners in chemokine regulation? Immunol. Cell Biol. 89, 197–206.

Horuk, R. (2015). The Duffy Antigen Receptor for Chemokines DARC/ACKR1. Front. Immunol. 6.

Huang, J., Rauscher, S., Nawrocki, G., Ran, T., Feig, M., de Groot, B.L., Grubmüller, H., and MacKerell, A.D. (2017). CHARMM36m: an improved force field for folded and intrinsically disordered proteins. Nat. Methods 14, 71–73.

Jia, R., Martens, C., Shekhar, M., Pant, S., Pellowe, G.A., Lau, A.M., Findlay, H.E., Harris, N.J., Tajkhorshid, E., Booth, P.J., et al. (2020). Hydrogen-deuterium exchange mass spectrometry captures distinct dynamics upon substrate and inhibitor binding to a transporter. Nat. Commun. 11.

Kaneko, J., and Kamio, Y. (2004). Bacterial Two-component and Hetero-heptameric Pore-forming Cytolytic Toxins: Structures, Pore-forming Mechanism, and Organization of the Genes. Biosci. Biotechnol. Biochem. 68, 981–1003.

Keener, J.E., Zhang, G., and Marty, M.T. (2020). Native Mass Spectrometry of Membrane Proteins. Anal. Chem. acs.analchem.0c04342.

Kong, C., Neoh, H., and Nathan, S. (2016). Targeting Staphylococcus aureus Toxins: A Potential form of Anti-Virulence Therapy. Toxins 8, 72.

Kufareva, I., Gustavsson, M., Zheng, Y., Stephens, B.S., and Handel, T.M. (2017). What Do Structures Tell Us About Chemokine Receptor Function and Antagonism? Annu. Rev. Biophys. 46, 175–198.

Latorraca, N.R., Venkatakrishnan, A.J., and Dror, R.O. (2017). GPCR Dynamics: Structures in Motion. Chem Rev 17.

Lau, A.M., Claesen, J., Hansen, K., and Politis, A. (2020). Deuteros 2.0: peptide-level significance testing of data from hydrogen deuterium exchange mass spectrometry. Bioinformatics btaa 677.

Lee, J., Cheng, X., Swails, J.M., Yeom, M.S., Eastman, P.K., Lemkul, J.A., Wei, S., Buckner, J., Jeong, J.C., Qi, Y., et al. (2016). CHARMM-GUI Input Generator for NAMD, GROMACS, AMBER, OpenMM, and CHARMM/OpenMM Simulations Using the CHARMM36 Additive Force Field. J. Chem. Theory Comput. 12, 405–413.

Los, F.C.O., Randis, T.M., Aroian, R.V., and Ratner, A.J. (2013). Role of Pore-Forming Toxins in Bacterial Infectious Diseases. Microbiol. Mol. Biol. Rev. 77, 173–207.

Lubkin, A., Lee, W.L., Alonzo, F., Wang, C., Aligo, J., Keller, M., Girgis, N.M., Reyes-Robles, T., Chan, R., O’Malley, A., et al. (2019). Staphylococcus aureus Leukocidins Target Endothelial DARC to Cause Lethality in Mice. Cell Host Microbe 25, 463-470.e9.

Mahoney, J.P., and Sunahara, R.K. (2016). Mechanistic insights into GPCR-G protein interactions. Curr. Opin. Struct. Biol. 41, 247–254.

Martens, C., and Politis, A. (2020). A glimpse into the molecular mechanism of integral membrane proteins through hydrogen-deuterium exchange mass spectrometry. Protein Sci.

Marty, M.T., Baldwin, A.J., Marklund, E.G., Hochberg, G.K.A., Benesch, J.L.P., and Robinson, C.V. (2015). Bayesian Deconvolution of Mass and Ion Mobility Spectra: From Binary Interactions to Polydisperse Ensembles. Anal. Chem. 87, 4370–4376.

Maurel, D., Comps-Agrar, L., Brock, C., Rives, M.-L., Bourrier, E., Ayoub, M.A., Bazin, H., Tinel, N., Durroux, T., Prézeau, L., et al. (2008). Cell-surface protein-protein interaction analysis with time-resolved FRET and snap-tag technologies: application to GPCR oligomerization. Nat. Methods 5, 561– 567.

Miszta, P., Pasznik, P., Jakowiecki, J., Sztyler, A., Latek, D., and Filipek, S. (2018). GPCRM: a homology modeling web service with triple membrane-fitted quality assessment of GPCR models. Nucleic Acids Res. 46, W387–W395.

Möller, I.R., Slivacka, M., Nielsen, A.K., Rasmussen, S.G.F., Gether, U., Loland, C.J., and Rand, K.D. (2019). Conformational dynamics of the human serotonin transporter during substrate and drug binding. Nat. Commun. 10.

Nariya, H., and Kamio, Y. (1997). Identification of the Minimum Segment Essential for the H γ ll-Specific Function of Staphylococcal γ -Hemolysin. Biosci. Biotechnol. Biochem. 61, 1786–1788.

Nibbs, R.J.B., and Graham, G.J. (2013). Immune regulation by atypical chemokine receptors. Nat. Rev. Immunol. 13, 815–829.

Novitzky-Basso, I., and Rot, A. (2012). Duffy antigen receptor for chemokines and its involvement in patterning and control of inflammatory chemokines. Front. Immunol. 3.

Ozawa, T., Kaneko, J., and Kamio, Y. (1995). Essential Binding of LukF of Staphylococcal γ -Hemolysin Followed by the Binding of H γ II for the Hemolysis of Human Erythrocytes. Biosci. Biotechnol. Biochem. 59, 1181–1183.

Peng, Z., Takeshita, M., Shibata, N., Tada, H., Tanaka, Y., and Kaneko, J. (2018). Rim domain loops of staphylococcal β-pore forming bi-component toxin S-components recognize target human erythrocytes in a coordinated manner. J. Biochem. (Tokyo) 164, 93–102.

Pronk, S., Páll, S., Schulz, R., Larsson, P., Bjelkmar, P., Apostolov, R., Shirts, M.R., Smith, J.C., Kasson, P.M., van der Spoel, D., et al. (2013). GROMACS 4.5: a high-throughput and highly parallel open source molecular simulation toolkit. Bioinformatics 29, 845–854.

Pruenster, M., Mudde, L., Bombosi, P., Dimitrova, S., Zsak, M., Middleton, J., Richmond, A., Graham, G.J., Segerer, S., Nibbs, R.J.B., et al. (2009). The Duffy antigen receptor for chemokines transports chemokines and supports their promigratory activity. Nat. Immunol. 10, 101–108.

Reading, E., Ahdash, Z., Fais, C., Ricci, V., Wang-Kan, X., Grimsey, E., Stone, J., Malloci, G., Lau, A.M., Findlay, H., et al. (2020). Perturbed structural dynamics underlie inhibition and altered efflux of the multidrug resistance pump AcrB. Nat. Commun. 11.

Reyes-Robles, T., Alonzo, F., Kozhaya, L., Lacy, D.B., Unutmaz, D., and Torres, V.J. (2013). Staphylococcus aureus Leukotoxin ED Targets the Chemokine Receptors CXCR1 and CXCR2 to Kill Leukocytes and Promote Infection. Cell Host Microbe 14, 453–459.

Seilie, E.S., and Bubeck Wardenburg, J. (2017). Staphylococcus aureus pore-forming toxins: The interface of pathogen and host complexity. Semin. Cell Dev. Biol. 72, 101–116.

Spaan, A.N., Vrieling, M., Wallet, P., Badiou, C., Reyes-Robles, T., Ohneck, E.A., Benito, Y., de Haas, C.J.C., Day, C.J., Jennings, M.P., et al. (2014). The staphylococcal toxins γ-haemolysin AB and CB differentially target phagocytes by employing specific chemokine receptors. Nat. Commun. 5.

Spaan, A.N., Reyes-Robles, T., Badiou, C., Cochet, S., Boguslawski, K.M., Yoong, P., Day, C.J., de Haas, C.J.C., van Kessel, K.P.M., Vandenesch, F., et al. (2015). Staphylococcus aureus Targets the Duffy Antigen Receptor for Chemokines (DARC) to Lyse Erythrocytes. Cell Host Microbe 18, 363–370.

Spaan, A.N., van Strijp, J.A.G., and Torres, V.J. (2017). Leukocidins: staphylococcal bi-component pore-forming toxins find their receptors. Nat. Rev. Microbiol. 15, 435–447.

Tromp, A.T., and van Strijp, J.A.G. (2020). Studying Staphylococcal Leukocidins: A Challenging Endeavor. Front. Microbiol. 11.

Tromp, A.T., Van Gent, M., Abrial, P., Martin, A., Jansen, J.P., De Haas, C.J.C., Van Kessel, K.P.M., Bardoel, B.W., Kruse, E., Bourdonnay, E., et al. (2018). Human CD45 is an F-component-specific receptor for the staphylococcal toxin Panton–Valentine leukocidin. Nat. Microbiol. 3, 708–717.

Turner, N.A., Sharma-Kuinkel, B.K., Maskarinec, S.A., Eichenberger, E.M., Shah, P.P., Carugati, M., Holland, T.L., and Fowler, V.G. (2019). Methicillin-resistant Staphylococcus aureus: an overview of basic and clinical research. Nat. Rev. Microbiol. 17, 203–218.

Vacchini, A., Locati, M., and Borroni, E.M. (2016). Overview and potential unifying themes of the atypical chemokine receptor family. J. Leukoc. Biol. 99, 883–892.

Vandenesch, F., Lina, G., and Henry, T. (2012). Staphylococcus aureus Hemolysins, bi-component Leukocidins, and Cytolytic Peptides: A Redundant Arsenal of Membrane-Damaging Virulence Factors? Front. Cell. Infect. Microbiol. 2.

Vasquez, M.T., Lubkin, A., Reyes-Robles, T., Day, C.J., Lacey, K.A., Jennings, M.P., and Torres, V.J. (2020). Identification of a domain critical for Staphylococcus aureus LukED receptor targeting and lysis of erythrocytes. J. Biol. Chem. 295, 17241–17250.

Vauquelin, G., and Charlton, S.J. (2013). Exploring avidity: understanding the potential gains in functional affinity and target residence time of bivalent and heterobivalent ligands: Exploring bivalent ligand binding properties. Br. J. Pharmacol. 168, 1771–1785.

Yamashita, D., Sugawara, T., Takeshita, M., Kaneko, J., Kamio, Y., Tanaka, I., Tanaka, Y., and Yao, M. (2014). Molecular basis of transmembrane beta-barrel formation of staphylococcal pore-forming toxins. Nat. Commun. 5.

Yamashita, K., Kawai, Y., Tanaka, Y., Hirano, N., Kaneko, J., Tomita, N., Ohta, M., Kamio, Y., Yao, M., and Tanaka, I. (2011). Crystal structure of the octameric pore of staphylococcal -hemolysin reveals the -barrel pore formation mechanism by two components. Proc. Natl. Acad. Sci. 108, 17314–17319.

Yen, H.-Y., Hopper, J.T.S., Liko, I., Allison, T.M., Zhu, Y., Wang, D., Stegmann, M., Mohammed, S., Wu, B., and Robinson, C.V. (2017). Ligand binding to a G protein–coupled receptor captured in a mass spectrometer. Sci. Adv. 3, e1701016.

Yen, H.-Y., Hoi, K.K., Liko, I., Hedger, G., Horrell, M.R., Song, W., Wu, D., Heine, P., Warne, T., Lee, Y., et al. (2018). PtdIns(4,5)P2 stabilizes active states of GPCRs and enhances selectivity of G-protein coupling. Nature 559, 423–427.

Yoong, P., and Torres, V.J. (2015). Counter inhibition between leukotoxins attenuates Staphylococcus aureus virulence. Nat. Commun. 6.

Zhang, H., Han, G.W., Batyuk, A., Ishchenko, A., White, K.L., Patel, N., Sadybekov, A., Zamlynny, B., Rudd, M.T., Hollenstein, K., et al. (2017). Structural basis for selectivity and diversity in angiotensin II receptors. Nature 544, 327–332.

Zhang, J., Yang, J., Jang, R., and Zhang, Y. (2015). GPCR-I-TASSER: A Hybrid Approach to G Protein-Coupled Receptor Structure Modeling and the Application to the Human Genome. Structure 23, 1538–1549.

Zhang, X., Chien, E.Y.T., Chalmers, M.J., Pascal, B.D., Gatchalian, J., Stevens, R.C., and Griffin, P.R. (2010). Dynamics of the β _2_ -Adrenergic G-Protein Coupled Receptor Revealed by Hydrogen−Deuterium Exchange. Anal. Chem. 82, 1100–1108.

Zheng, J., Strutzenberg, T., Pascal, B.D., and Griffin, P.R. (2019). Protein dynamics and conformational changes explored by hydrogen/deuterium exchange mass spectrometry. Curr. Opin. Struct. Biol. 58, 305–313.

Zwier, J.M., Roux, T., Cottet, M., Durroux, T., Douzon, S., Bdioui, S., Gregor, N., Bourrier, E., Oueslati, N., Nicolas, L., et al. (2010). A Fluorescent Ligand-Binding Alternative Using Tag-lite ® Technology. J. Biomol. Screen. 15, 1248–1259.

