## Supplemental Information for "Mechanisms of GPCR hijacking by *Staphylococcus aureus*"

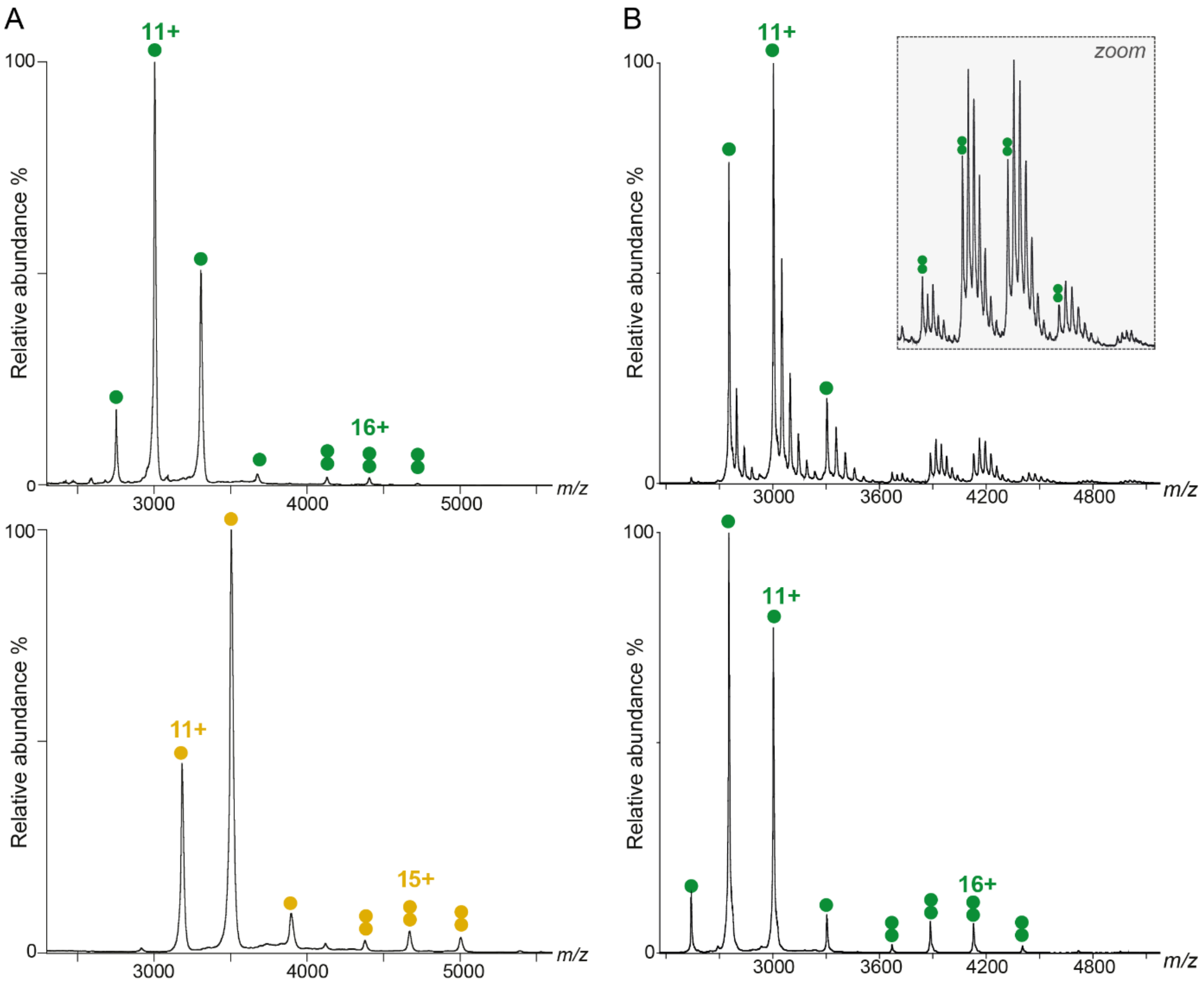


**Supplementary Figure S1. Native MS spectra of HlgA and HlgB.** A) nMS spectrum of 10 μM HlgA (Top) and HlgB (bottom), showing the presence of both monomeric and dimeric species in ammonium acetate buffer in the absence of detergent. B) nMS spectrum of 28 μM HlgA in ammonium acetate buffer in the presence of 2CMC DDM detergent, at low (top) and high (bottom) activation energies. Detergent-binding is visible for both monomeric and dimeric HlgA species, and DDM molecules can be dissociated as we higher the activation in the collision cell. Green circles HlgA (monomeric 33 004 ± 1 Da and dimeric 66 061 ± 14 Da) and yellow circles HlgB (monomeric 34 943 ± 1 Da and dimeric 69 972 ± 10 Da).

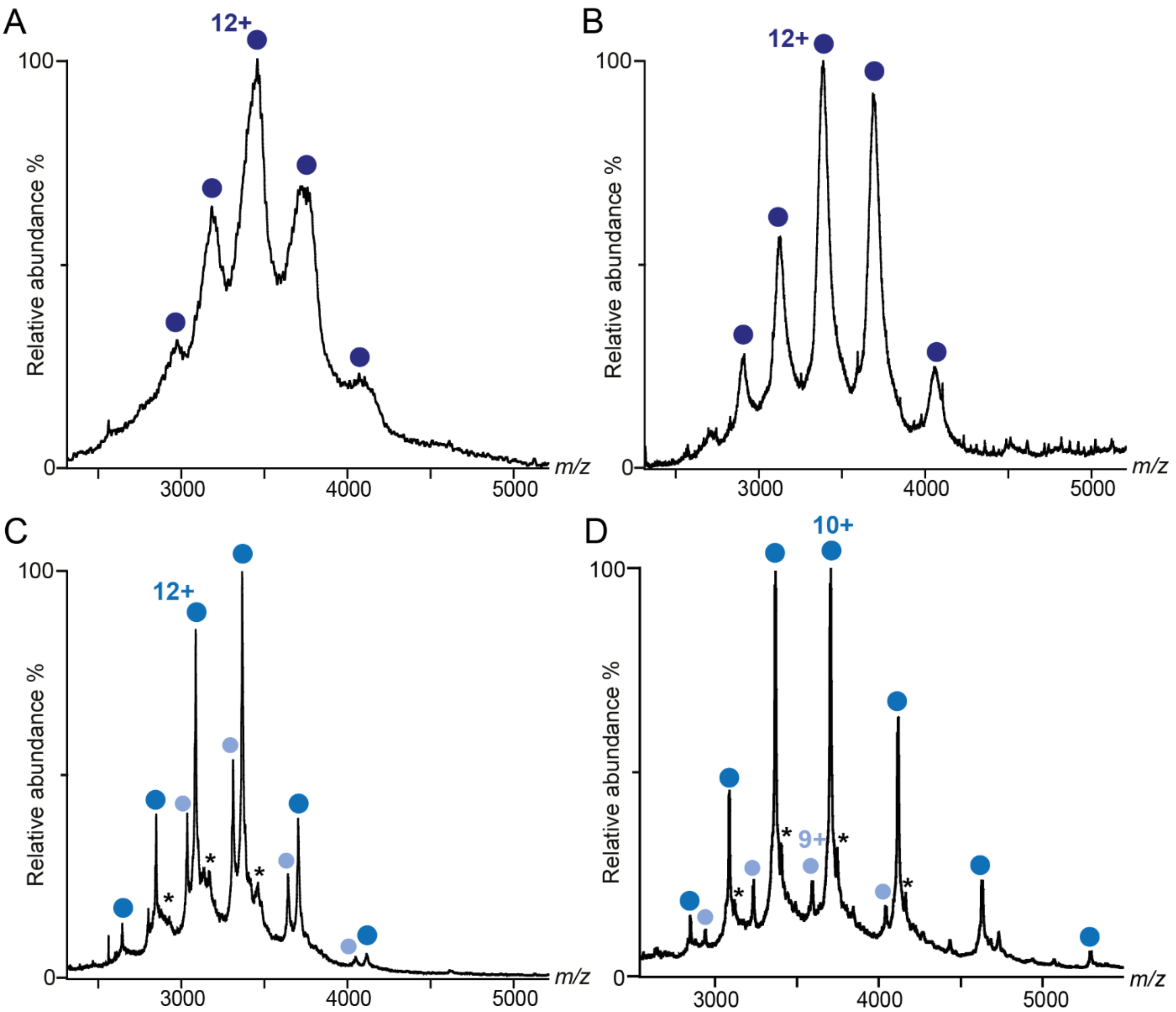


**Supplementary Figure S2. Native MS spectra of different monomeric ACKR1 productions.** A) ACKR1 WT produced in SF9 cells, showing large unresolved peaks between 40 and 42 kDa, as opposed to the theoretical mass of 37 kDa for unmodified ACKR1. B) ACKR1 WT produced in HEK GnTI$^{-}$ cells, showing less broad peaks due to more homogeneous glycosylations, however the measure mass of 40 531 ± 32 Da was still higher than the expected one. C) ACKR1 WT treated with PNGase F, showing the main isoform at 37 022 ± 1 Da (blue circles) and a Cter-hydrolysed form at 36 411 ± 1 Da (light blue circle). Additional isoforms are also present at lower relative intensities (asterisk). D) N$^{16,27,33}$Q-ACKR1 produced in SF9 cells showing the main isoform at 37 005 ± 1 Da (blue circles) and an Nter-hydrolysed form at 32 304 ± 1 Da (light blue circle). Additional isoforms are also present at lower relative intensities (asterisk).

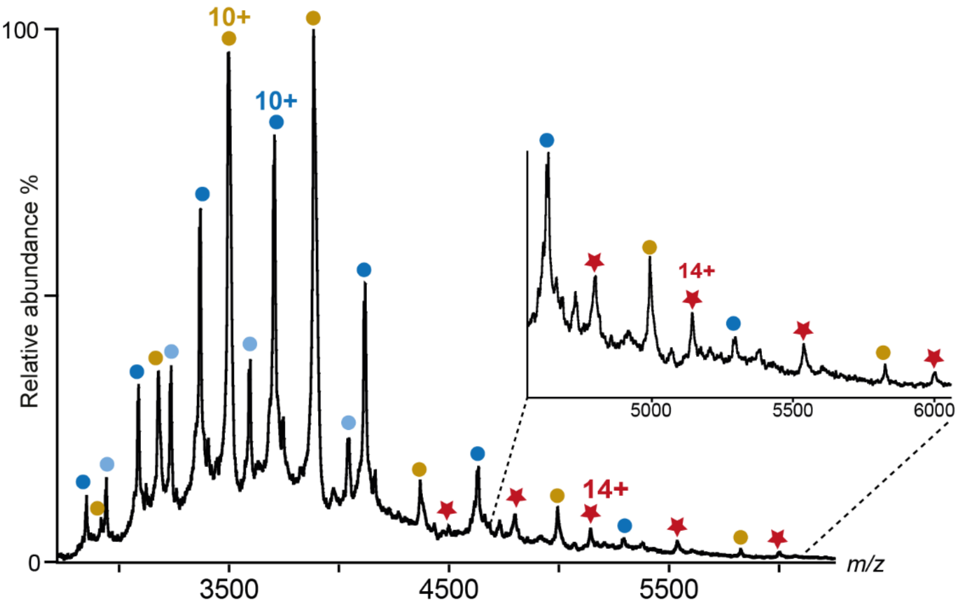


**Supplementary Figure S3. Native MS spectrum of HlgB–ACKR1 complex**. nMS spectrum of a mixture of 2 μM HlgB and 5 μM N$^{16,27,33}$Q-ACKR1 deglycosylated mutant, showing the presence of monomeric HlgB (yellow circles, 34 943 ± 1 Da), N$^{16,27,33}$Q-ACKR1 (blue circles, 37 005 ± 1 Da) and partially-hydrolysed N$^{16,27,33}$Q-ACKR1 (light blue circles, 32 304 ± 1 Da). Complexes formed between HlgB and full-length N$^{16,27,33}$Q-ACKR1 are labelled with dark red stars (71 993 ± 29 Da). No complexes were visible between HlgB and the hydrolysed form of N$^{16,27,33}$Q-ACKR1, implying that the hydrolysis of ACKR1 observed is at the Nter part of the receptor, which correlates with the different masses and might implicate a role of glycosylations in the *in vitro* stabilisation of the relatively long and unstructured Nter part of ACKR1.

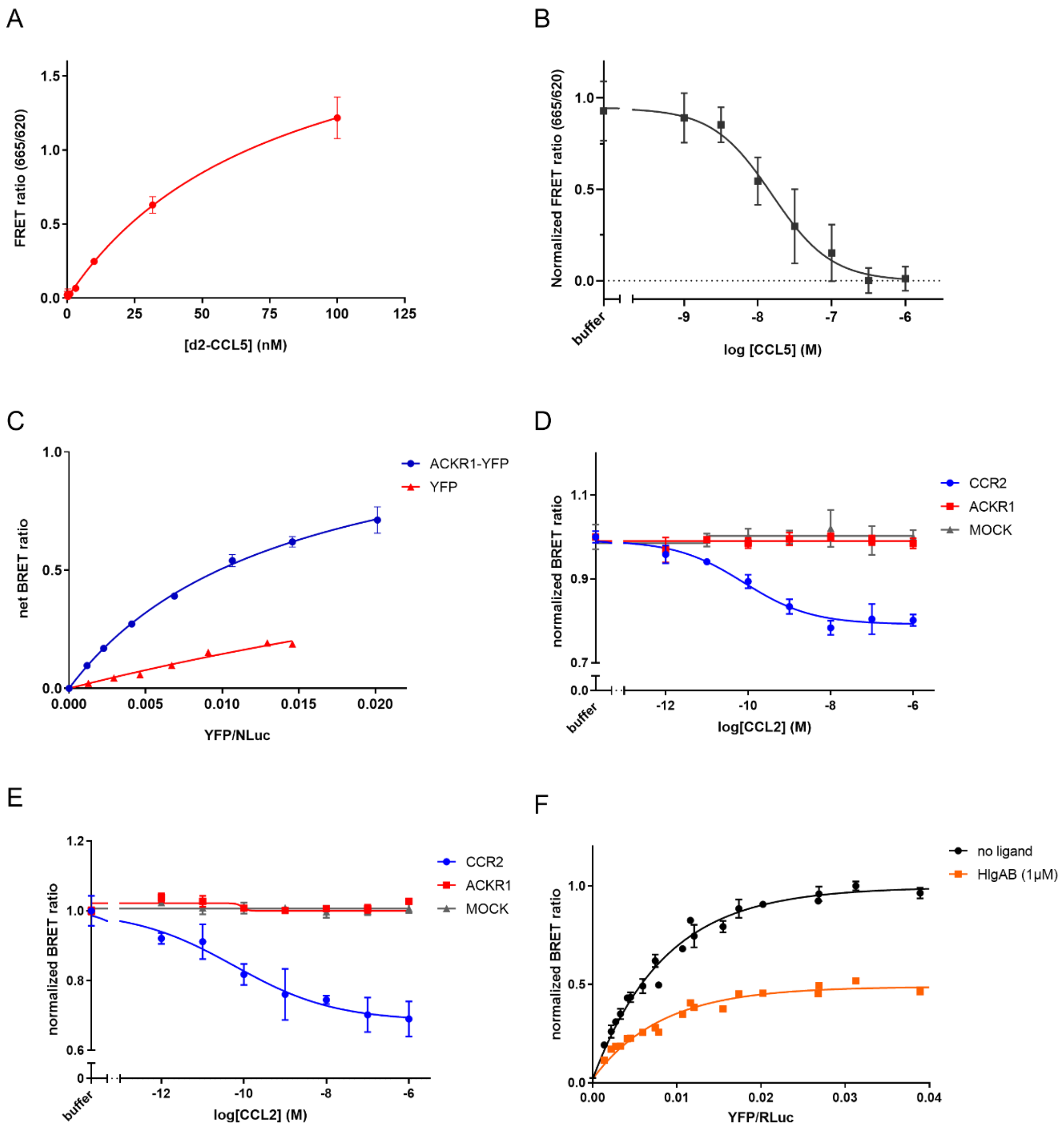


**Supplementary Figure S4**. **Cell-based assays supplementary data.** A) Saturation curve showing TR-FRET ratio between SNAP-ACKR1 labelled with Tb and d2-CCL5 upon addition of increasing amounts of d2-CCL5. A $K_d$ value of 78 ± 13 nM was obtained. Data shown are the mean +/- SEM of one experiment performed in triplicates. B) Competitive binding showing the dose-response decrease in TR-FRET ratio upon binding of unlabelled CCL5 to SNAP-ACKR1 labelled with Tb and d2-CCL5. An IC50 of 16 nM corresponding to a Ki ~ 14 nM (with d2-CCL5 concentration = 12 nM) was obtained. Data shown are the mean +/- SEM of two independent experiments performed in triplicates. C) ACKR1-YFP and ACKR1-NLuc BRET saturation upon increasing the concentration of ACKR1-YFP but not YFP alone. Data shown are the mean +/- SEM of one experiment performed in triplicates and are representative of 3 independent experiments. D) and E) BRET assays between $G\alpha$-RLuc and $\beta\delta$-YFP to follow G-protein activation showing activation of Goa (D) and of Gi1 (E) by CCR2 but not by ACKR1 upon addition of CCL2 ligand. Data shown are the mean +/- SEM of one experiment performed in triplicates and are representative of two independent experiments. Similar results show no activation in the presence of ACKR1 with CCL5 ligand. F) BRET saturation curve between

ACKR1-YFP and G$\alpha$i1-RLuc showing the interaction between ACKR1 C-terminal part and G$\alpha$i1 in living cells, and specific decreased BRET signal upon addition of HlgAB regardless the used YFP: RLuc ratio. Data shown are the mean +/- SEM of 3 independent experiments performed in triplicates.

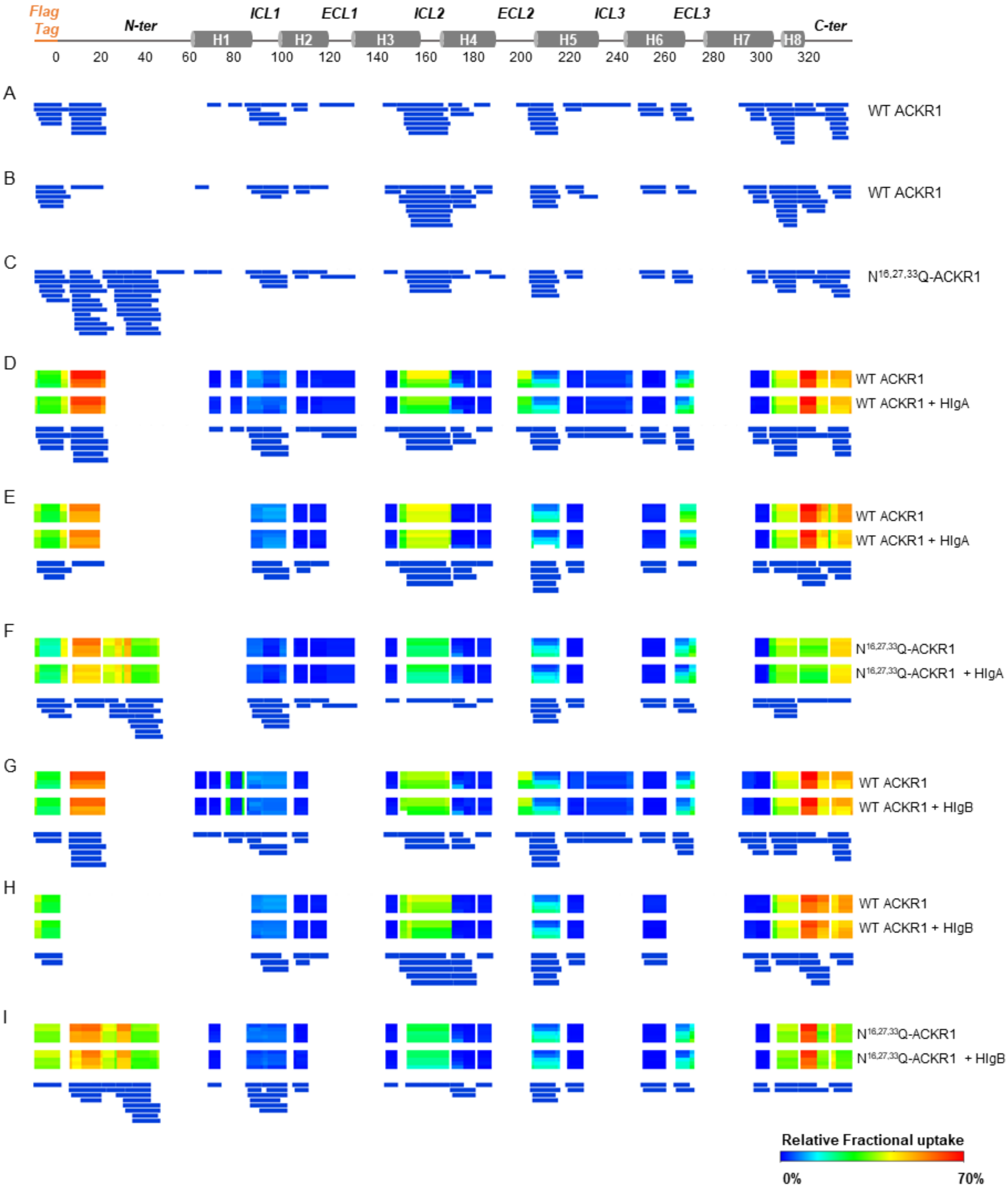


**Supplementary Figure S5. Coverage maps and relative fractional uptakes of ACKR1 biological replicates.** A) and B) Coverage maps of both biological replicates of WT ACKR1 and C) of one replicate of deglycosylated $N^{16,27,33}$Q-ACKR1 where the N-terminal region of ACKR1 is more covered, all in the absence of added leukotoxins. 77 peptides, 70.1% coverage and 3.41 redundancy (A); 66 peptides, 59.5% coverage and 3.42 redundancy (B); 92 peptides, 72.7% coverage and 4.06 redundancy (C). Each blue bar represents one detected and manually validated peptide, and relate to the scheme representing the various theoretical transmembrane helices and loops with their corresponding residue numbers based on our generated ACKR1 model. D) to E) Relative fractional uptakes and common coverage maps for differential HDX analysis of all biological replicates analysed with WT and $N^{16,27,33}$Q-ACKR1 in the presence of HlgA and HlgB. For additional information, see supported documents (xls files).

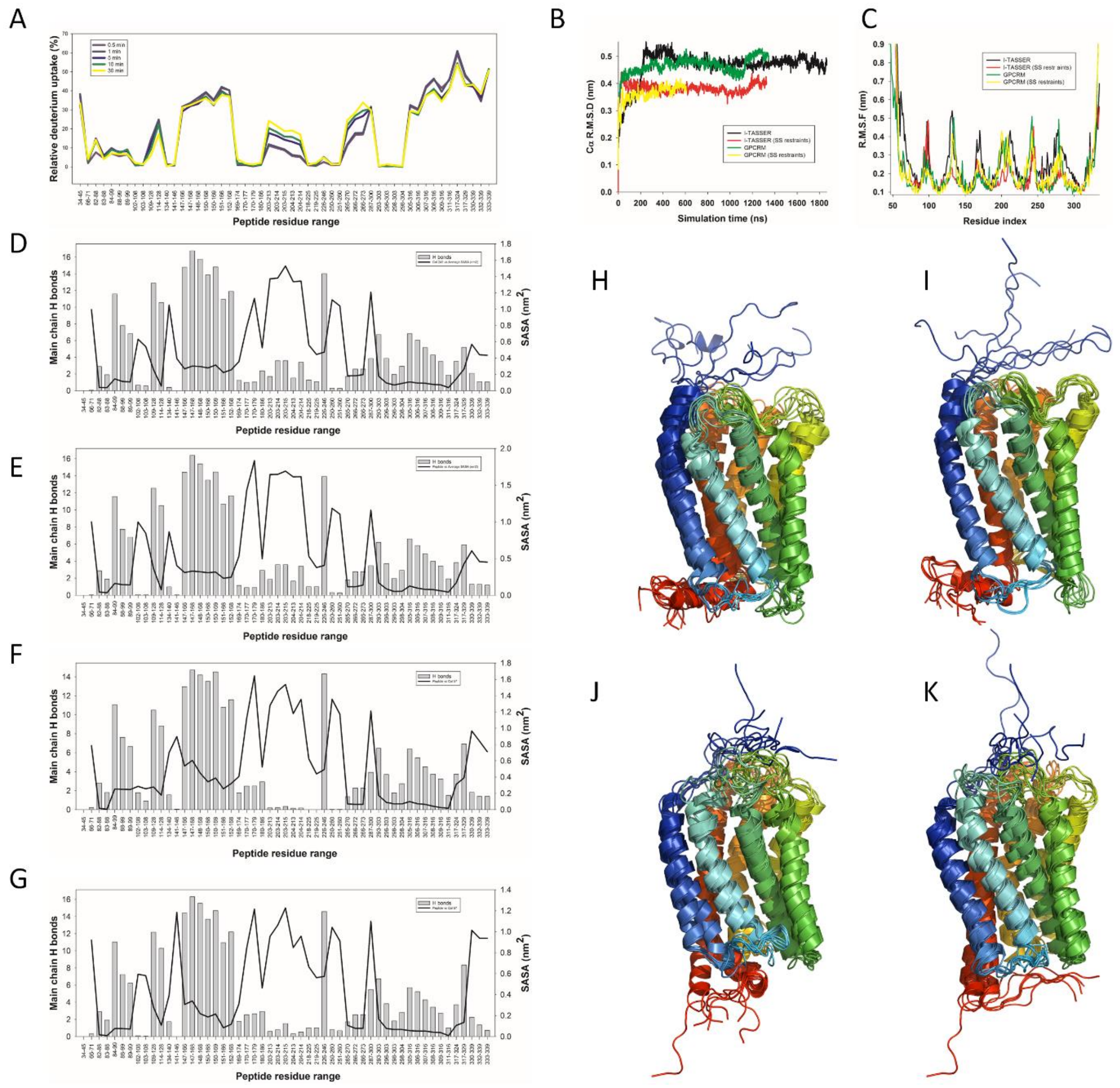


**Supplementary Figure S6. Analysis of molecular dynamics simulations and correlation with ACKR1 HDX data.** A) Relative fractional uptake of deuterium for ACKR1 peptides. B) Root mean square deviation (RMSD) of Cα atoms from the starting structure for residues 51-320 of the I-TASSER and GPCRM models in the presence or absence of SS restraints. C) Root mean square fluctuations (RMSF) of Cα atoms along the protein sequence. D), E), F), and G) Main chain SASA and hydrogen bond profiles extracted from MDS trajectories of unrestrained I-TASSER model (D), restrained I-TASSER model (E), unrestrained GPCRM model (F) and restrained GPCRM model (G). The number of hydrogen bonds is shown as a grey bar, and the average SASA is shown as a black line for the experimentally characterized ACKR1 peptides. H), I), J) and K) Superimposed snapshots of ACKR1 models extracted from MDS trajectories of unrestrained I-TASSER model (H), restrained I-TASSER model (I), unrestrained GPCRM model (J) and restrained GPCRM model (K). For each trajectory, seven models corresponding to t=0, 100 … 600 ns are shown in cartoon representation and colored from blue (N-terminus) to red (C-terminus).
